# An aging-sensitive compensatory secretory phospholipase that confers neuroprotection and cognitive resilience

**DOI:** 10.1101/2024.07.26.605338

**Authors:** Cinzia Vicidomini, Travis D. Goode, Kathleen M. McAvoy, Ruilin Yu, Conor H. Beveridge, Sanjay N. Iyer, Matheus B. Victor, Noelle Leary, Liam Evans, Michael J. Steinbaugh, Zon Weng Lai, Marina C. Lyon, Manuel Rico F.S Silvestre, Gracia Bonilla, Ruslan I. Sadreyev, Tobias C. Walther, Shannan Ho Sui, Takaomi Saido, Kei Yamamoto, Makoto Murakami, Li-Huei Tsai, Gaurav Chopra, Amar Sahay

**Affiliations:** Center for Regenerative Medicine, Massachusetts General Hospital, Boston, Massachusetts, USA; Harvard Stem Cell Institute, Cambridge, Massachusetts, USA; Department of Psychiatry, Massachusetts General Hospital, Harvard Medical School, Boston, Massachusetts, USA; BROAD Institute of MIT and Harvard, Cambridge, Massachusetts, USA; Department of Chemistry, Purdue University, West Lafayette, IN 47907, USA; Picower Institute for Learning and Memory, Massachusetts Institute of Technology, Cambridge, Massachusetts, USA; Department of Brain and Cognitive Sciences, Massachusetts Institute of Technology, Cambridge, Massachusetts, USA; Harvard Chan Bioinformatics Core, Harvard School of Public Health, Harvard University, Boston, Massachusetts, USA; Harvard Chan Advanced Multi-omics Platform, Harvard T.H. Chan School of Public Health, Boston, Massachusetts, USA; Department of Molecular Metabolism, Harvard T.H. Chan School of Public Health, Boston, Massachusetts, USA; Department of Cell Biology, Harvard Medical School, Boston, Massachusetts, USA; Department of Molecular Biology. Massachusetts General Hospital, Harvard Medical School, Boston, Massachusetts, USA; Howard Hughes Medical Institute, Boston, Massachusetts, USA; Laboratory for Proteolytic Neuroscience, RIKEN Center for Brain Science, Saitama 351-0198, Japan; Graduate School of Technology, Industrial and Social Sciences, Tokushima University, 2-1 Minami-jyosanjima, Tokushima 770-8513, Japan; Laboratory of Microenvironmental and Metabolic Health Sciences, Center for Disease Biology and Integrative Medicine, Graduate School of Medicine, The University of Tokyo, 7-3-1 Hongo, Bunkyo-ku, Tokyo 113-8655, Japan; Purdue Institute for Integrative Neuroscience, Purdue University, West Lafayette, IN 47907, USA; Purdue Institute for Drug Discovery, Purdue University, West Lafayette, IN 47907, USA; Purdue Center for Cancer Research, Purdue University, West Lafayette, IN 47907, USA; Regenstrief Center for Healthcare Engineering, Purdue University, West Lafayette, IN 47907, USA

## Abstract

Breakdown of lipid homeostasis is thought to contribute to pathological aging, the largest risk factor for neurodegenerative disorders such as Alzheimer’s Disease (AD). Cognitive reserve theory posits a role for compensatory mechanisms in the aging brain in preserving neuronal circuit functions, staving off cognitive decline, and mitigating risk for AD. However, the identities of such mechanisms have remained elusive. A screen for hippocampal dentate granule cell (DGC) synapse loss-induced factors identified a secreted phospholipase, *Pla2g2f*, whose expression increases in DGCs during aging. *Pla2g2f* deletion in DGCs exacerbates aging-associated pathophysiological changes including synapse loss, inflammatory microglia, reactive astrogliosis, impaired neurogenesis, lipid dysregulation and hippocampal-dependent memory loss. Conversely, boosting *Pla2g2f* in DGCs during aging is sufficient to preserve synapses, reduce inflammatory microglia and reactive gliosis, prevent hippocampal-dependent memory impairment and modify trajectory of cognitive decline. Ex vivo, neuronal-PLA2G2F mediates intercellular signaling to decrease lipid droplet burden in microglia. Boosting *Pla2g2f* expression in DGCs of an aging-sensitive AD model reduces amyloid load and improves memory. Our findings implicate PLA2G2F as a compensatory neuroprotective factor that maintains lipid homeostasis to counteract aging-associated cognitive decline.

## Main

The hippocampus plays a crucial role in registration of our experiences as episodic memories and includes the dentate gyrus (DG) subregion that contributes to memory formation by decreasing memory interference ^1–4^. Furthermore, dysfunction in DG is considered to be a primary driver of aging-associated cognitive decline and a hallmark of mild cognitive impairment ^5–9^. Healthy cognitive aging depends on a balance of resilience and vulnerability factors that dictate the tempo of pathophysiological alterations, such as synapse loss ^10–12^, lipid dysregulation ^13,14,15–18^, emergence of maladaptive “inflammatory” microglia ^15,19–24^ and reactive astrogliosis ^25–27^, oxidative stress ^28–30^, loss of hippocampal neurogenesis ^31–37^, and excitation-inhibition imbalance ^7,9,38,39,7,9,40^. Disruption of this balance leads to a trajectory of cognitive decline and increased risk for mild cognitive impairment and Alzheimer’s disease (AD).

A complementary perspective, first offered within the framework of cognitive reserve theory, posits that pathological cognitive decline reflects a failure to compensate for the progressive increase in pathophysiological alterations over time (**Fig. 1a**)^41^. Studies in rodents and humans suggest a role for re-organization of neural activity patterns and network level functional connectivity, increase in large diameter head dendritic spines, and changes in synaptic gene expression as substrates for functional compensation during aging ^42–48^; however, the identities of the underlying mechanisms have remained elusive. Consequently, and in contrast to our understanding of how genetic factors mediating resilience and vulnerability influence cognitive reserve in aging ^41,49–55^, a mechanistic instantiation of how compensation confers cognitive resilience in aging is still lacking. Evidence for endogenous compensatory cognitive resilience mechanisms should fulfill the following criteria. First, such a mechanism must be naturally engaged in response to pathophysiological alterations during aging. Second, blockade or disruption of such a mechanism must exacerbate aging-associated pathophysiological alterations. Third, boosting such a mechanism must confer cognitive resilience and prevent cognitive decline during aging.

**Fig. 1:**
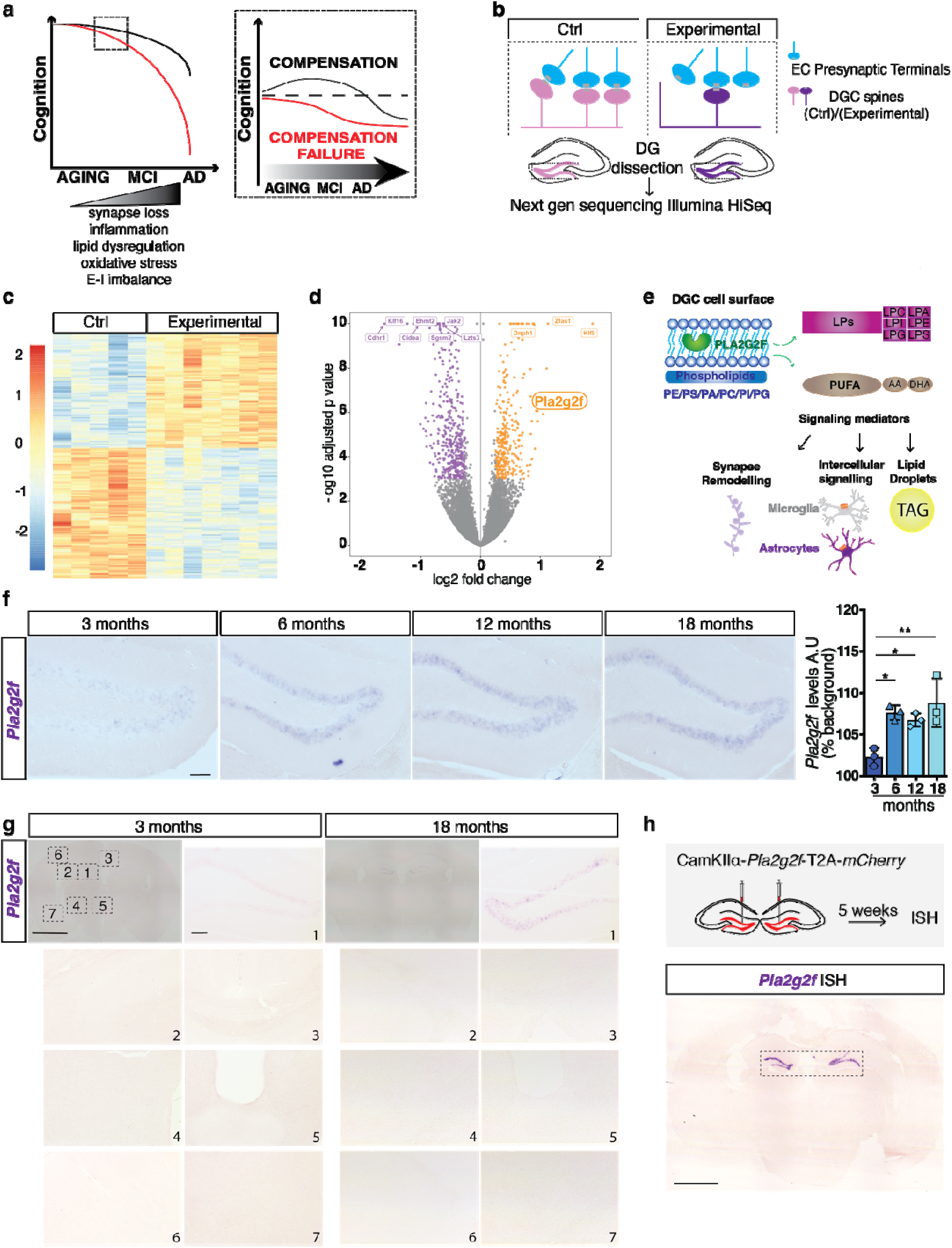
Pla2g2f is upregulated in DGCs following synapse loss and during aging. **a**, Schematic conveying concept of how loss of endogenous compensatory mechanisms during aging increases risk for mild cognitive impairment (MCI) and Alzheimer’s Disease (AD). **b**, Schematic of genetic screen to identify DGC synapse-loss induced compensatory factors. The dentate gyrus of Ctrl and Experimental groups were dissected and processed for Illumina next-generation sequencing (n=5 Ctrl, n=7 Experimental). **c**, Heatmap showing hierarchical clustering of differentially expressed genes (DEGs). Row-wise scaled Z score expression values are shown, ranging from high (orange) to low (blue) expression +/- 3 standard deviations (n=5 Ctrl, n=7 Experimental). **d**, Volcano plot of statistical significance (−log10 adjusted p value) vs. magnitude of change (log2 of fold change) of gene expression. Upregulated genes in experimental mice are on the right and downregulated genes in experimental mice are on the left (entire dataset for differentially expressed genes reported in Supplementary Tables 1 and 2). *Pla2g2f* is upregulated in the experimental group. **e**, Schematic illustrating PLA2G2F dependent enzymatic hydrolysis of cell surface glycerophospholipids (PE, Phosphatidylethanolamine /PS, Phosphatidylserine /PA, Phosphatidic acid/PC, Phosphatidylcholine /PI, Phosphatidylinositol /PG, Phosphatidylglycerol) into lysophospholipids (LPE/LPS/LPA/LPC/LPI/LPG) and PUFAs (polyunsaturated fatty acids, eg: AA (arachidonic acid) and DHA (Docosahexaenoic acid) to mediate synapse remodeling, intercellular communication and lipid droplet regulation. **f**, Representative images and quantification of *Pla2g2f* expression (in situ hybridization) in the dentate gyrus subregion of hippocampus of C57Bl/6J mice (3, 6, 12,18 months old) (n=3 animals per group; scale bar 100μm). **g**, Representative images of *Pla2g2f* expression (in situ hybridization) in the brain of C57Bl/6J mice (3 and 18 months old mice). Insets 1,2,3,4,5,6,7 show higher magnification images of different brain regions: 1 (dentate gyrus), 2 (CA2/3), 3 (corpus callosum and overlying cingulate cortex), 4 (dorsal medial hypothalamic nucleus), 5 (paraventricular thalamic nucleus), 6 (primary somatosensory cortex), 7 (entopeduncular nucleus) (scale bar whole brain panels 250μm; scale bar different brain regions 100μm). **h**, Validation of *Pla2g2f* riboprobe and viral vector to boost *Pla2g2f* expression in DG. Representative image showing *Pla2g2f* mRNA levels in the dentate gyrus, 5 weeks after lentiviral injection (scale bar 250μm). Data are displayed as mean ± SEM; \*\**P*<0.005, \**P*<0.05 [Ordinary one-way ANOVA, Dunnett’s multiple comparisons test in f].

Secreted phospholipases comprise a family of highly conserved Ca^2+^-dependent enzymes ^56^ that exhibit cell-type specific regulation of expression that is sensitive to aging ^57^, pathogen infection ^57^, oxidative stress ^57^, lipopolysaccharide treatment ^58^, ferroptosis^59^, high fat diet-induced inflammation ^60^ and ischemia ^61^. Secreted phospholipase-mediated hydrolysis of membrane glycerophospholipids into lysophospholipids and polyunsaturated fatty acids directly influences membrane lipid composition, membrane fluidity, synaptic remodeling, vesicle membrane fusion and exocytosis ^62–65^. Additionally, membrane-derived lysophospholipids and polyunsaturated fatty acids signal through cell-surface receptors and lipid metabolites to mediate cell-cell communication and regulate a plethora of biological processes ^56,60,61,66,67^. Thus, depending on cellular context (i.e membrane lipid composition and cell-type diversity in the milieu), secreted phospholipases exert both cell-autonomous and non-cell autonomous mechanisms to protect against, compensate for or potentiate impairments triggered by homeostatic insults.

In an *in vivo* genetic screen designed to identify synapse-loss induced factors in the DG, we found that expression of a secreted phospholipase, *Pla2g2f* ^58,67^, was upregulated in dentate granule cells (DGCs) following synapse loss. *Pla2g2f* expression is undetectable in the adult brain, but its expression increases in DGCs during aging. Inducible deletion of *Pla2g2f* in DGCs in middle-aged mice exacerbated aging-associated loss of synapses, emergence of inflammatory microglia (defined by expression of lysosomal protein CD68 and hypo-ramified morphology)^15,19–22^ and reactive astrogliosis, disruption of lipid homeostasis and hippocampal-dependent memory impairment. Boosting *Pla2g2f* in DGCs of aged mice prevented these pathophysiological impairments and preserved hippocampal-dependent memory. Additionally, elevating *Pla2g2f* expression levels in DGCs of middle-aged mice modified trajectory of aging-associated cognitive decline. Using human microglia-neuronal conditioned medium coculture experiments, we showed that PLA2G2F mediates intercellular signaling to decrease lipid burden in microglia. Boosting *Pla2g2f* expression in an aging-sensitive AD mouse model reduced amyloid burden and preserved hippocampal-dependent memory. Our data implicate PLA2G2F as a compensatory neuroprotective and resilience factor that maintains cognitive reserve by preserving healthy brain aging and counteracting cognitive decline.

## Results

### *Pla2g2f* expression is upregulated in DGCs following synapse loss and during aging

To identify factors that mediate or moderate loss of synapses, we designed an *in vivo* functional screen to profile gene expression changes in DGCs following inducible loss of perforant path-DGC synapses. Prior work from our laboratory identified Kruppel-like factor 9 (KLF9) as a negative regulator of dendritic spines^31,68,69^. We previously engineered and characterized an inducible genetic knock-in mouse model in which we can bidirectionally regulate *Klf9* levels and the number of PSD95 positive dendritic spines in DGCs ^31^. Using this genetic system, we performed next-generation sequencing of RNA isolated from DGCs following synapse loss (**Fig. 1b**). Annotation of gene expression revealed downregulation of modules involved in neuronal remodeling and upregulation of ribosomal biogenesis and translation at synapses (**Extended Data Fig. 1**). Analysis of differentially expressed genes identified a secreted phospholipase, *Pla2g2f*, as significantly upregulated (1.93 fold, p=9X ^10–7^)(**Fig. 1c,d, Supplementary Tables 1,2**). *Pla2g2f* encodes for a type II secreted phospholipase A2 (sPLA_2_-IIF) which hydrolyzes the fatty acyl chain on the sn2 position in 3-sn-phosphoglycerides. Unlike many other members of the secreted PLA_2_ family, the functional importance of PLA2G2F has not been established in the central nervous system and the highest known expression of *Pla2g2f* is in keratinocytes^67^. Prior work had shown that inflammation (Lipopolysaccharide or LPS injection) induces *Pla2g2f* expression in the brain and peripheral tissues ^58^. Based on documented role of secreted phospholipases in regulation of lipid metabolism, intercellular communication, lipid remodelling and inflammation^56,60,61,67^, we hypothesized that PLA2G2F-dependent hydrolysis of glycerophospholipids in DGCs and generation of lysophospholipids and bioactive signaling mediators following synapse loss may promote synapse remodeling, neuron-glia/microglia intercellular communication, and restore lipid homeostasis (**Fig. 1e**). As a first step towards testing this hypothesis, we mapped *Pla2g2f* expression in the brain by *in situ* hybridization. We found that *Pla2g2f* expression is negligible in the adult brain but is gradually upregulated in a sparse population of DGCs during aging when perforant path-DGC synapses are lost (**Fig. 1f,g**). We validated the riboprobe used in this analysis by assessing *Pla2g2f* expression after viral mediated boosting of *Pla2g2f* expression in DGCs (**Fig. 1h**). Based on the expression pattern of *Pla2g2f* alone, it is not possible to determine if elevation in *Pla2g2f* expression during aging mediates vulnerability or compensatory resilience to aging-associated pathophysiological alterations in the hippocampus and cognitive decline. To distinguish between these possibilities, we deployed genetic and viral recombination systems to selectively ablate *Pla2g2f* expression in DGCs during aging.

### Loss of *Pla2g2f* in DGCs exacerbates aging-associated pathophysiological and hippocampal-dependent memory impairment

Perforant path-DGC synapses are lost during aging^10^. To determine how loss of *Pla2g2f* in DGCs affects synapses, we induced recombination of *Pla2g2f* in 16 months old mice that were bigenic for CamKIIα−Cre^ERT2^ transgene and *Pla2g2f* conditional alleles and stereotactically injected low-titer virus expressing *mCherry* into the dentate gyrus (**Fig. 2a**). Tamoxifen dependent Cre^ERT2^ recombination of *Pla2g2f* conditional alleles in CamKIIα−Cre^ERT2^-*Pla2g2f ^f/f^* mice abolishes *Pla2g2f* expression in DGCs (**Extended Data Fig. 2a,b**). Analysis of dendritic spines of sparsely labeled DGCs in these mice at 20 months of age showed a reduction in dendritic spine density following loss of *Pla2g2f* (**Fig. 2b**, **Extended Data Fig. 2c,d**). Quantification of spine head diameters revealed a reduction in large head diameter spines suggesting an increase in synapse loss ^70^(**Fig. 2b**).

**Fig. 2:**
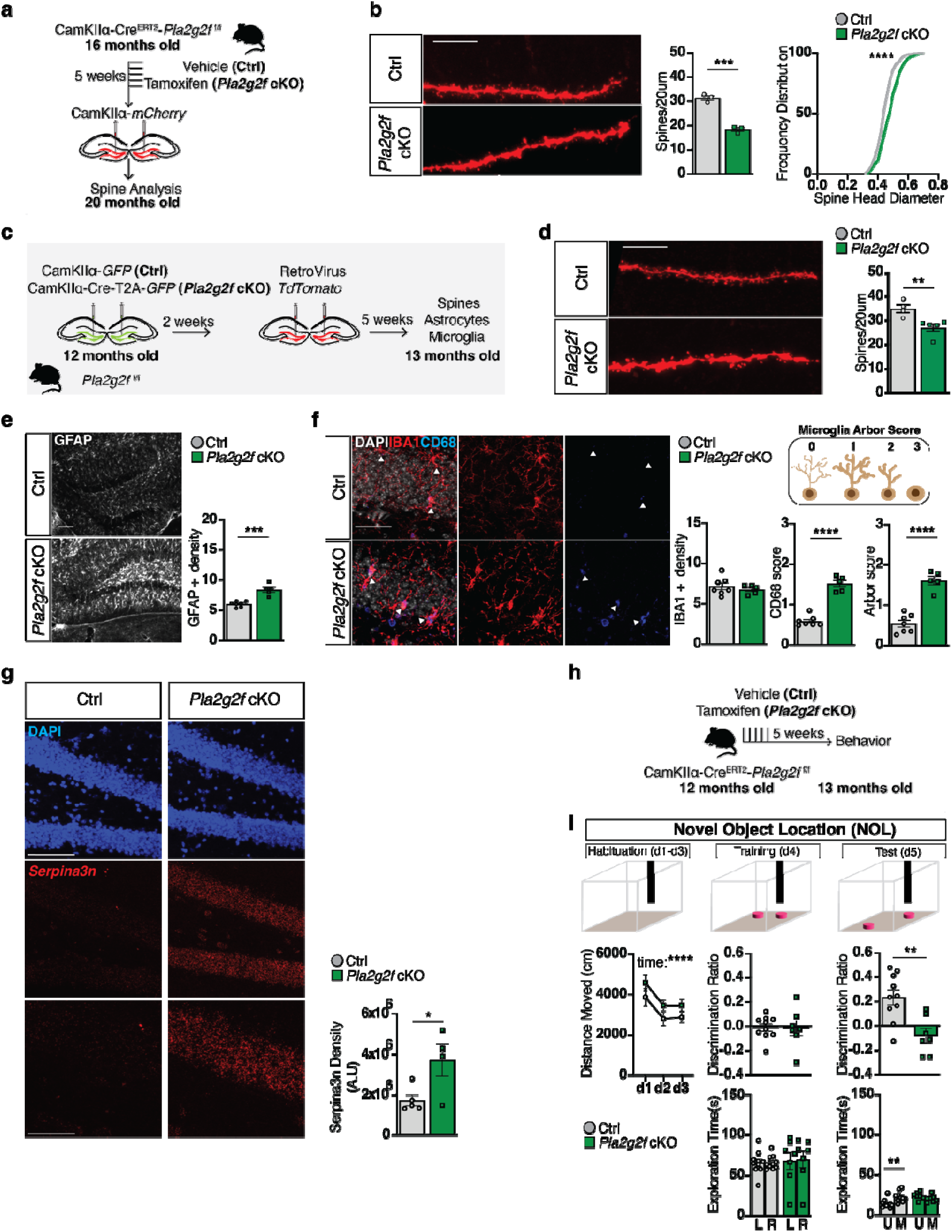
Loss of *Pla2g2f* in middle-age promotes dendritic spine loss, inflammatory microglia, reactive astrocytes, and hippocampal-dependent memory impairment. **a**, CamKIIα-Cre^ERT2^-*Pla2g2f ^f/f^* bigenic mice (16 months old) were injected with vehicle (Ctrl) or tamoxifen (referred to as *Pla2g2f* cKO) for 5 days; 5 weeks later, mice were injected with CamKIIα-*mCherry* lentiviruses to label dendritic spines of DGCs and dendritic spine analysis was performed 4 months later in 20 months old mice. **b**, Representative images and quantification of dendritic spine density and spine head diameter frequency distribution of dendritic spines of DGCs of 20 months old mice (n= 3 mice per group; scale bar, 10μm). **c**, Experimental design showing lentiviral injections to express CamKIIα-GFP or Cre-T2A-GFP in DGCs of 12 months old *Pla2g2f ^f/f^* mice followed by injection of low titer retroviruses expressing tdTomato to sparsely label newly generated DGCs. Analysis of dendritic spine density of adult-born DGCs, astrocytes, and microglia was performed 5 weeks after retroviral injection in 13 months old mice. **d**, Representative images and quantification of dendritic spine density of 5-week-old adult-born DGCs in the dentate gyrus of 13 months old mice (n= 4 Ctrl and n=5 *Pla2g2f* cKO; scale bar, 10μm). **e**, Representative images and quantification of GFAP positive astrocyte density in the dentate gyrus of 13 months old Ctrl and *Pla2g2f* cKO mice (n=7 Ctrl, n=5 *Pla2g2f* cKO; scale bar, 100μm). Y axis represents the average of GFAP + cells per region of interest (ROI) per mouse. **f**, Representative images and quantification of IBA1 positive microglia density, CD68 score, and arbor score of microglia in the dentate gyrus of Ctrl and *Pla2g2f cKO* (13 months old mice). Box contains schematic showing the microglia arbor score criteria used for the analysis (n=7 Ctrl, n=5 *Pla2g2f* cKO; arrowheads point to individual microglia cells; scale bar, 100μm). DAPI, 4′,6-diamidino-2-phenylindole. Y axis represents the average of IBA1+ cell per ROI per mouse. **g**, Representative images and quantification of *Serpina3n* expression in the dentate gyrus of 13 months old Ctrl and *Pla2g2f* cKO mice (n= 5Ctrl, n=4 *Pla2g2f* cKO; scale bar 100μm, scale bar higher magnification bottom panels 50μm). **h**, CamKIIα-Cre^ERT2^-*Pla2g2f ^f/f^*bigenic mice were injected with vehicle or tamoxifen for 5 days and 5 weeks later spatial learning and memory was assessed. **i**, Hippocampal dependent spatial learning and memory was assessed in the novel object location test. Discrimination Ratio at training represents: (exploration time of left (L) object - exploration time of right (R) object) / (exploration time of left object + exploration time of right object). Discrimination Ratio at test represents: (exploration time of the moved (M) object - exploration time of unmoved (U) object)/(exploration time of the moved object + exploration time of unmoved object) (n=10 Ctrl, n=7 *Pla2g2f* cKO). Data are displayed as mean ± SEM; \*\*\*\**P*<0.0001, \*\*\**P*=0.0002, \*\**P*<0.005, \**P*<0.05 [unpaired t-test in b(left panel), d, e, f, g, i (middle and right top panels); repeated measures two way ANOVA in i (top left panel, middle and right bottom panels) Sidak’s multiple comparison test in i (right bottom panel); Kolmogorov-Smirnov test in B(right panel)].

We next asked whether PLA2G2F in DGCs of middle-aged mice affects maturation of adult-born DGCs. We injected lentiviruses expressing CamKIIα-GFP or Cre-T2A-GFP into DG of 12 months old *Pla2g2f ^f/f^* (*Pla2g2f cKO*) mice and then two weeks later injected low titer retroviruses expressing tdTomato to sparsely label newly generated DGCs (**Fig. 2c,d**, **Extended Data Fig. 2e,f**). Analysis of dendritic spines of tdTomato-labeled maturing DGCs (5 weeks old neurons) showed a reduction in dendritic spine density (**Fig. 2d**). Analysis of mossy fiber terminals of newly generated DGCs did not reveal a difference in mossy fiber terminal size, number of filopodia per mossy fiber terminal, or parvalbumin inhibitory neuron synapses in hippocampal area CA3 suggesting that PLA2G2F does not regulate output connectivity of DGCs^39,71^(**Extended Data Fig. 3a-c**). Together, these results demonstrate that PLA2GF functions non-cell autonomously or in a paracrine manner to regulate dendritic spines of DGCs and that PLA2G2F is necessary for adult hippocampal neurogenesis, specifically, maturation of adult-born DGCs in middle-aged mice.

**Fig. 3:**
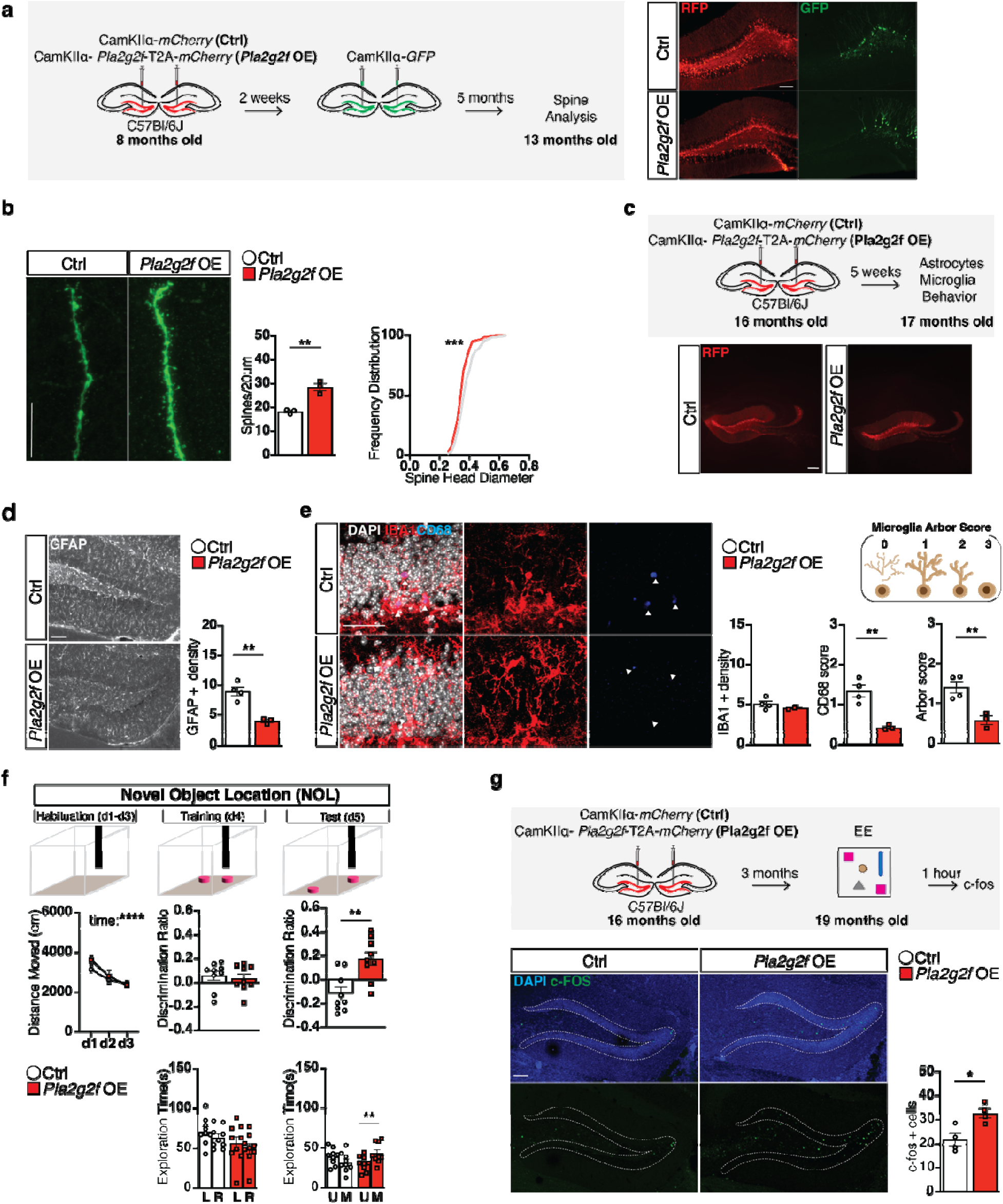
Boosting *Pla2g2f* in aging reverses impairments in dendritic spines, microglia, astrocytes and hippocampal-dependent memory. **a**, CamKIIα-*mCherry* (Ctrl) or CamKIIα-*Pla2g2f*-T2A-*mCherry* (*Pla2g2f* OE) expressing lentiviruses were injected into the dentate gyrus of 8 months old C57BL/6J mice followed by injection of low titer CamKIIα-*GFP* lentiviruses to sparsely label dendritic spines of DGCs. 5 months later spine analysis was performed in13 months old mice, Left panel. Representative images of dentate gyrus of 13 months old mice expressing the two lentiviral expression systems, Right panel. **b**, Representative images and quantification of dendritic spine density and spine head diameter frequency distribution of dendritic spines of DGCs of 13 month old mice (n= 3 mice per group; scale bar, 10μm). **c**, Top panel: experimental design to boost *Pla2g2f* expression in DGCs of 16 months old C57Bl/6J mice. Analysis of astrocytes, microglia, and spatial learning and memory was performed 5 weeks later in17 months old mice (scale bar, 100μm). Bottom panel: representative images of 17 months old mice dentate gyrus expressing CamKIIα-*mCherry* (Ctrl) or CamKIIα-*Pla2g2f*-T2A-*mCherry* (*Pla2g2f* OE) lentiviruses (scale bar 250μm). **d**, Representative images and quantification of GFAP positive astrocytes in the dentate gyrus of 17 months old Ctrl and *Pla2g2f* OE mice (n=4 Ctrl, n=3 *Pla2g2f* OE; scale bar, 100μm). Y axis represents the average of GFAP + cell per ROI per mouse. **e**, Representative images and quantification of IBA1 positive density, CD68 score, and arbor score of microglia in the dentate gyrus of 17 months old Ctrl and *Pla2g2f* OE mice. Box contains schematic showing the microglia arbor score criteria used for the analysis (n=4 Ctrl, n=3 *Pla2g2f* OE; arrowheads point to individual microglia cells; scale bar, 100μm). DAPI, 4′,6-diamidino-2-phenylindole. Y axis represents the average of IBA1+ cell per ROI per mouse. **f**, Hippocampal dependent spatial learning and memory was assessed in the novel object location test. Discrimination Ratio at training represents: (exploration time of left (L) object - exploration time of right (R) object) / (exploration time of left object + exploration time of right object). Discrimination Ratio at test represents: (exploration time of the moved (M) object - exploration time of unmoved (U) object)/(exploration time of the moved object + exploration time of unmoved object) (n=9 Ctrl, n=9 *Pla2g2f* OE). **g**, Top panel: experimental design to boost *Pla2g2f* expression in DGCs of 16 months old C57Bl/6J mice. 3 months later (19 months old) mice were placed in an enriched environment followed by c-FOS immunostaining. Bottom panel: Representative images and quantification of c-FOS positive neurons in the dentate gyrus of the hippocampus (n= 4 Ctrl, n=4 *Pla2g2f* OE; scale bar 100μm). Data are displayed as mean ± SEM; \*\*\*\**P*<0.0001,\*\*\**P*=0.0004, \*\**P*<0.005, \**P*<0.05 [unpaired t-test in b(left panel), d, e, g, f(middle and right top panels); repeated measures two way ANOVA in f (top left panel, middle and right bottom panels) Sidak’s multiple comparison test in f (right bottom panel); Kolmogorov-Smirnov test in b(right panel)].

Since secreted phospholipase-dependent hydrolysis of cell-surface glycerophospholipids generates bioactive lipid signaling mediators that may support intercellular communication, we asked how deletion of *Pla2g2f* in middle-aged mice affects the milieu of DGCs, and in particular, two hallmarks of aging: the emergence of inflammatory microglia (a cell state defined by increased phagocytosis, amoeboid morphology and higher levels of lysosomal marker CD68) and reactive astrocytes. We found that loss of *Pla2g2f* in DGCs increased the density of GFAP+ astrocytes, elevated CD68 levels in IBA1+ microglia, and transformed microglial arbor and morphology (less ramified and more amoeboid in shape) (**Fig. 2e,f**). Aging and neuroinflammation are associated with elevation in levels of the serine protease inhibitor, alpha 1-Antichymotrypsin or SERPINA3N ^26,27,72–74^. *Serpina3n* expression is increased in the brains of mice harboring *ApoE4* alleles, the strongest aging-associated risk factor for AD^75,76^, and SERPINA3N levels are associated with increased amyloid burden and inversely correlated with cognitive status ^74,77–80^. Deletion of *Pla2g2f* in DGCs resulted in elevation in *Serpina3n* expression in the dentate gyrus (**Fig. 2g**).

To understand how synapse loss, increased neuroinflammation and astrogliosis resultant from *Pla2g2f* deletion in middle-age affects hippocampal dependent memory, we tested a new cohort of CamKIIα−Cre^ERT2^-*Pla2g2f ^f/f^* mice in an aging-sensitive hippocampal-dependent task, novel object location ^81,82^, that is reliant on dentate gyrus dependent encoding of configural relationships between objects and their locations within contexts. This task probes a fundamental property of the hippocampus which is to bind object and spatial information into conjunctive representations or “contexts” ^83,84^. Middle-aged mice lacking *Pla2g2f* and control littermates exhibited comparable levels of habituation of the novel context on days 1-3 and comparable performance on training day 4. On test day, middle-aged mice lacking *Pla2g2f* spent equivalent amounts of time investigating the unmoved and displaced objects indicative of impaired hippocampal dependent memory. In contrast, controls spent more time investigating the displaced object, indicative of intact hippocampal dependent memory (**Fig. 2h,i**). Middle-aged mice lacking *Pla2g2f* exhibited normal anxiety-like behavior as assessed in the open field paradigm and elevated plus maze, and demonstrated normal cognition in the novel object recognition task that is less sensitive to dentate gyrus impairments (**Extended Data Fig. 4a-e**). Together, these results, suggest that loss of *Pla2g2f* in DGCs in middle-age impairs hippocampal dependent memory.

**Fig. 4:**
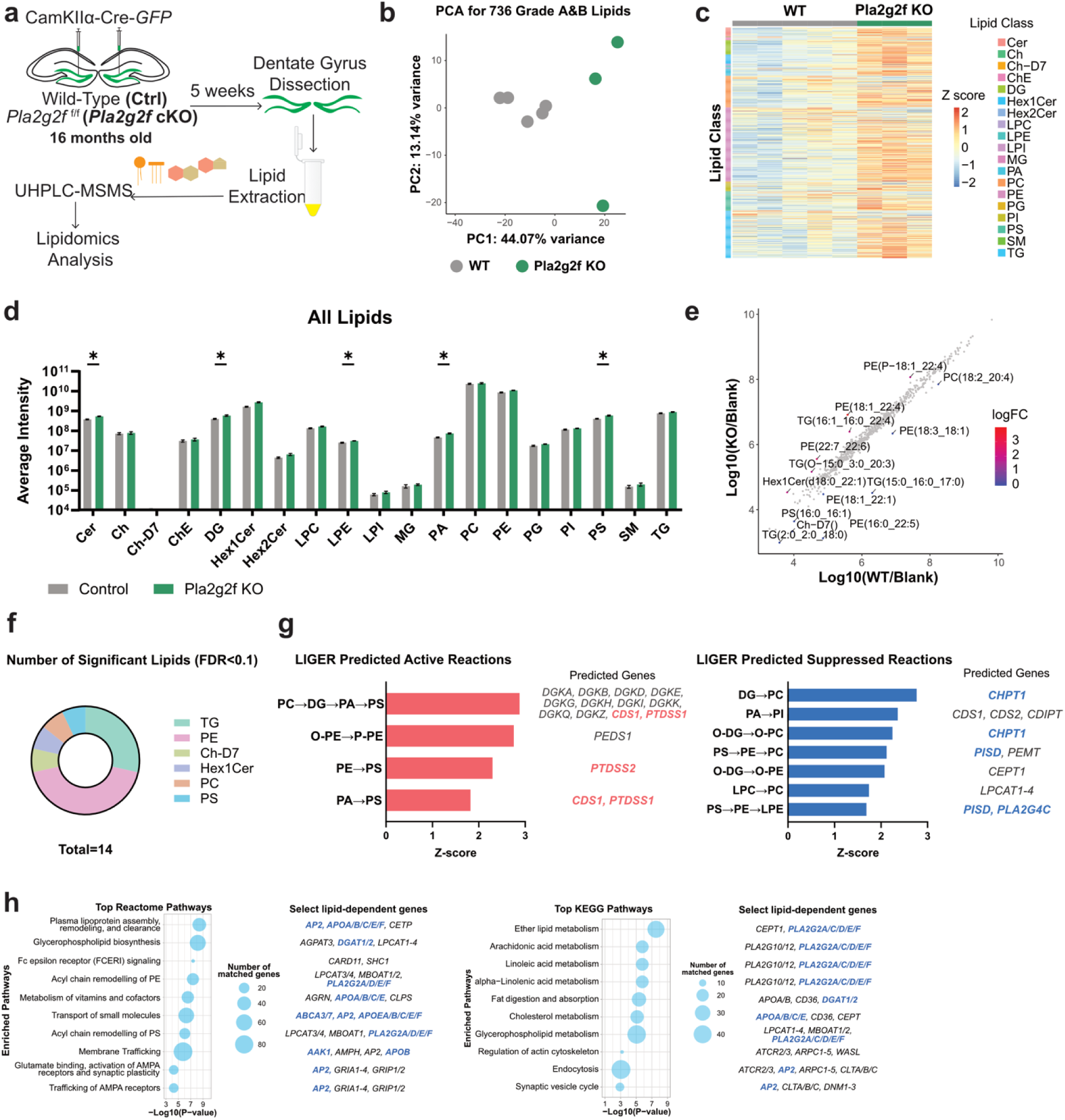
PLA2G2F is necessary for lipid homeostasis in aged dentate gyrus. **a**, Experimental design to virally recombine *Pla2g2f* in the dentate gyrus of 16 months old *Pla2g2f cKO* mice. N = 5 and N = 3 biological replicates were analyzed for control and *Pla2g2f* cKO, respectively. **b**, Principal component analysis (PCA) plot of lipid abundance for control (gray) and *Pla2g2f* cKO (green) samples. **c**, Heatmap for *Pla2g2f* cKO vs. control demonstrating changes in lipid abundance and showing Z-score. **d**, Average ion intensities of all lipids summed by class. Significance is calculated by unpaired multiple t-tests. * p-value < 0.05. **e**, Scatterplot showing abundance of individual lipids (grey dots) in WT control vs. *Pla2g2f* cKO animals. Lipids with FDR<0.1 are highlighted and labeled. **f**, Breakdown of significantly changed lipids by class (FDR<0.1). **g**, Lipid pathway enrichment analysis predicted active and suppressed lipid conversion reactions, and genes associated with reactions based on significantly different lipids (FDR < 0.1). Common genes which appeared in more than one lipid conversion reactions are highlighted. **h**, LipidSig pathway analysis based on significantly different lipids (FDR<0.1) using Reactome (left) or KEGG (right) database. The size of bubbles indicate number of lipid-dependent genes in the pathway. Select lipid-dependent genes are shown next to the corresponding pathways.

### Boosting *Pla2g2f* expression in DGCs in aging reverses pathophysiological and memory impairments and modifies trajectory of cognitive decline

To determine how boosting *Pla2g2f* expression in DGCs of middle-aged mice affects DGC synapses, we injected lentiviruses expressing CamKIIα−*mCherry* or CamKIIα−*Pla2g2f*-T2A-*mCherry* into the DG of 8 months old C57BL/6J mice. Two weeks later, we injected low titer lentivirus expressing CamKIIα-GFP and quantified dendritic spines of sparsely labeled DGCs 5 months later in 13 months old mice (**Fig. 3a**). Boosting *Pla2g2f* expression resulted in increased dendritic spine density and more large head diameter spines (**Fig. 3b**).

We next asked whether boosting *Pla2g2f* expression in DGCs protects against aging-associated neuroinflammation. We injected lentiviruses expressing CamKIIα−*mCherry* or CamKIIα−*Pla2g2f*-T2A-*mCherry* into the DG of 16 months old C57BL/6J mice and quantified microglia and astrocytes (**Fig. 3c**). Boosting *Pla2g2f* expression in aging mice engendered exactly the opposite changes in GFAP and microglia to that seen following loss of *Pla2g2f* in DGCs. Specifically, we found a decrease in the density of GFAP+ astrocytes in the dentate gyrus, reduction in CD68 levels in IBA1+ microglia, loss of amoeboid microglial shape and an increase in microglial arbor complexity (**Fig. 3d,e**).

Since boosting *Pla2g2f* expression in aged mice protects against synapse loss, neuroinflammation and reactive gliosis, we asked whether hippocampal-dependent memory is also preserved. We injected lentiviruses expressing CamKIIα−*mCherry* or CamKIIα−*Pla2g2f*-T2A-*mCherry* into the DG of a new cohort of 16 months old C57BL/6J mice and 5 weeks later we assessed hippocampal-dependent memory and anxiety-like behavior (**Fig. 3f**, **Extended Data Fig. 5**). While aged control mice exhibited impaired hippocampal-dependent memory and spent equivalent amounts of time investigating both the unmoved and displaced objects, aged mice with elevated *Pla2g2f* expression in DGCs exhibited intact hippocampal-dependent memory (**Figure 3f**), comparable to that seen in middle-aged controls (**Fig. 2h,i**). Boosting *Pla2g2f* expression did not affect anxiety-like behavior as assessed in the open field paradigm and elevated plus maze or performance in the novel object recognition task (**Extended Data Fig. 5a-e**). Consistent with enhanced DG function, initial exposure to a novel environment of open field elicited greater exploration in aged mice with elevated PLA2G2F levels in DGCs than controls (**Extended Data Fig. 5c**)^85,86^.

**Fig. 5:**
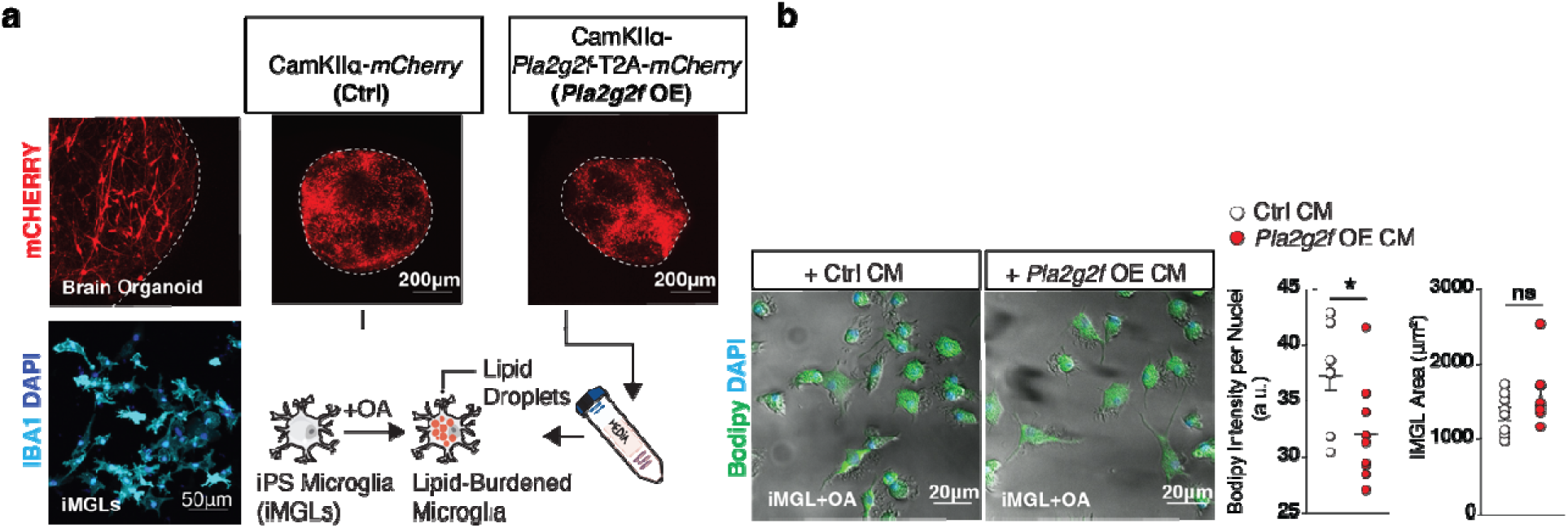
PLA2G2F mediates neuron-microglia signaling to reduce lipid burden. **a**, Excitatory neurons in brain organoids labeled with CamKIIα-*mCherry* (Ctrl) or CamKIIα-*Pla2g2f*-T2A-*mCherry* (*Pla2g2f* OE) lentiviruses at 120 days *in vitro*, Top left panel; microglia-like cells (iMGLs) derived in parallel from same donor iPSCs stained with the macrophage-specific marker, IBA, Bottom left panel (scale bar 50μm). 5-7 organoids were infected with either CamKIIα-*mCherry* (Ctrl) or CamKIIα-*Pla2g2f*-T2A-*mCherry* (*Pla2g2f* OE) for 18 days. Conditioned media from brain organoids was collected and applied to iMGLs that had been pre-treated with 3mM of Oleic Acid (OA), **b**, Middle panels. Representative images of OA treated microglia accumulating lipid droplets, labelled with the lipid neutral dye bodipy (in green). Images shown for iMGLs after 24 hours of incubation with either brain organoid conditioned media (CM) from Ctrl or *Pla2g2f* OE cultures (scale bar 20μm). Bodipy intensity was assessed following CM incubation for 24 hours to determine levels of intracellular lipid accumulation, Right bottom panels. iMGL morphology was not differentially impacted by treatment as shown by quantification of cell area (n= Average of 3 images for each of 9 independent cultures per group). Data are displayed as mean ± SEM; \**P*<0.05, n.s. not significant [Student’s t-test in b].

Since large diameter spines of DGCs are postsynaptic to perforant path inputs, we asked whether the PLA2G2F-dependent increase in large diameter spines of DGCs resulted in greater recruitment of the dentate gyrus of aged mice during exploration of a novel environment. Consistently, we found that boosting *Pla2g2f* expression in DGCs of aged mice resulted in more DGCs expressing the immediate early gene product c-FOS following exploration of a novel enriched environment (**Fig. 3g**).

If the increase in *Pla2g2f* expression in DGCs during aging confer cognitive resilience, then boosting *Pla2g2f* expression in middle-age should modify trajectory of cognitive decline during aging (**Extended Data Fig. 6a**). To test this hypothesis, we virally expressed CamKIIα−*mCherry* or CamKIIα−*Pla2g2f*-T2A-*mCherry* in DGCs of 12 months old C57BL/6J mice and probed cognitive function at 19 months of age (**Extended Data Fig. 6b,c**). Aged mice in which *Pla2g2f* expression was boosted in middle-age exhibited normal anxiety-like behavior (**Extended Data Fig. 6d**) but exhibited a strong trend towards enhanced hippocampal dependent memory in the novel object location task compared with the aged control group that was unable to cognitively perform the task (**Extended Data Fig. 6e**). A similar strong trend for cognitive enhancement was seen in another hippocampal dependent task, the pre-exposure-dependent contextual fear conditioning paradigm ^87^, in which mice have to retrieve and link a previously experienced context with a mild footshock presented to them in the same context (**Extended Data Fig. 6f**). Specifically, aged mice in which *Pla2g2f* expression was boosted in middle-age exhibited a strong trend towards higher levels of freezing behavior in the retrieval phase of testing compared to controls. Together, these results suggest that boosting *Pla2g2f* in DGCs during aging counteracts cognitive decline to confer cognitive resilience.

**Fig. 6:**
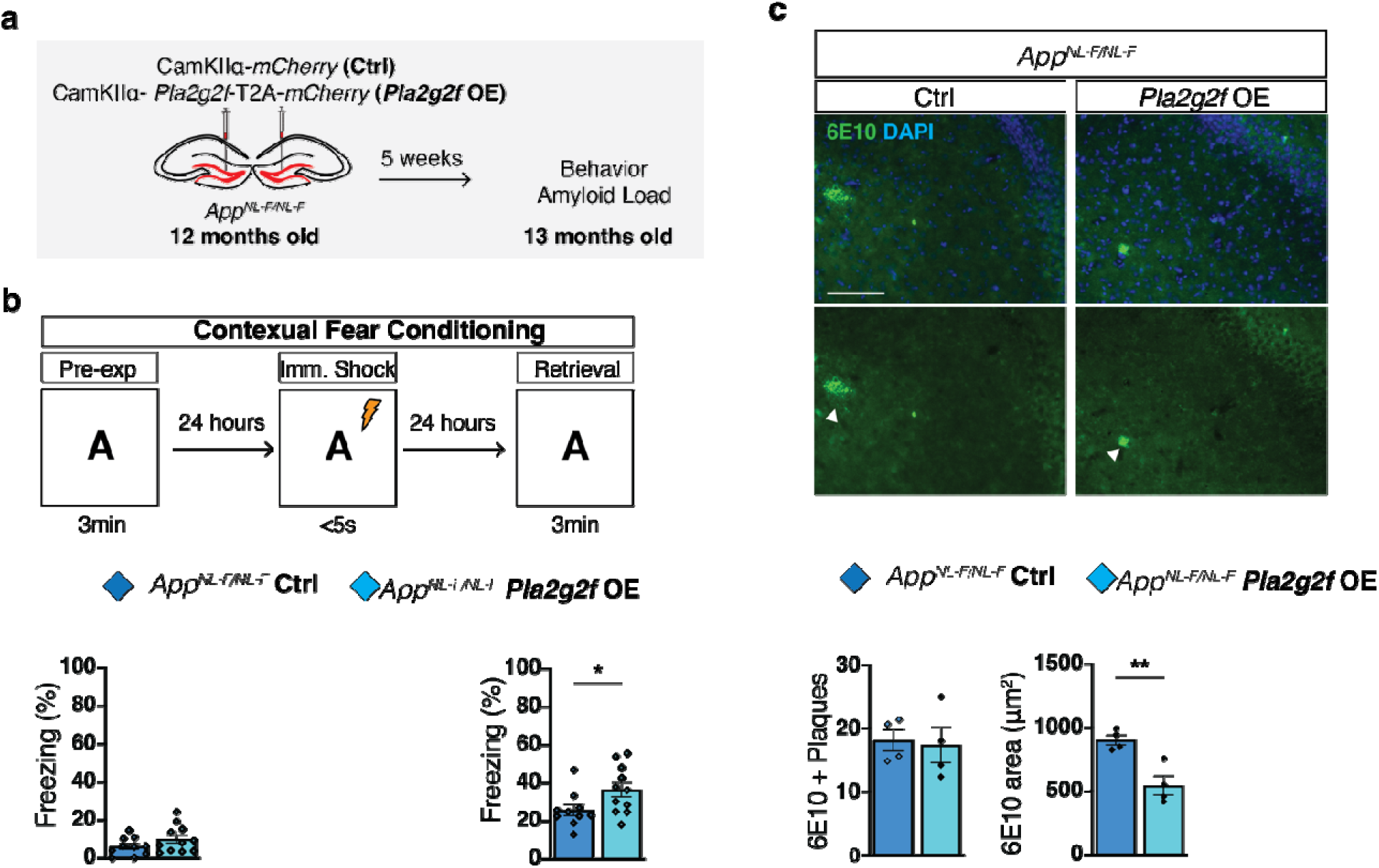
Boosting *Pla2g2f* in an aging-sensitive AD model preserves memory and decreases amyloid load. **a**, Experimental design to boost *Pla2g2f* expression in DGCs of 12 months old *App^NL-F/NL-F^* mice. Analysis of amyloid plaques and contextual fear memory was performed 5 weeks later in 13 months old mice. **b**, Contextual fear memory was assessed in pre-exposure-dependent contextual fear conditioning paradigm. Baseline freezing was assessed as the percentage of time spent freezing prior to fear conditioning (n=10 Ctrl, n=11 *Pla2g2f* OE). **c**, Representative images and quantification of 6E10 amyloid positive plaques and 6E10 amyloid plaque area in the hippocampus (n=4 Ctrl, n=4 *Pla2g2f* OE; scale bar 100μm). Data are displayed as mean ± SEM; \*\**P*<0.005, \**P*<0.05 [unpaired t-test in b, c].

### PLA2G2F is necessary for lipid homeostasis in aging and mediates neuronal-microglial signaling to reduce lipid burden

To understand how loss of PLA2G2F in DGCs of aged mice disrupts glycerophospholipid hydrolysis and lipid homeostasis, we performed lipidomic profiling and characterized lipid composition of the dentate gyrus following deletion of *Pla2g2f*. We injected CamKIIα−Cre-T2A-GFP viruses into the dentate gyrus of 16 months old *Pla2g2f* cKO mice and wild-type controls, dissected the dentate gyrus, extracted and separated lipid species using liquid chromatography and mass spectrometry (**Fig. 4**). Our lipidomics pipeline profiled a total number of 736 lipids among 19 classes. A majority of the lipid signal came from phosphatidylcholine (PC) and phosphatidylethanolamine (PE), the most abundant lipids in eukaryotic cells^88^(**Extended Data Fig. 7a, Supplementary Table 3**). Analysis of lipid abundance in absence of *Pla2g2f* revealed clear differences in lipid class and species profiles (**Fig. 4b,c**, **Extended Data Fig. 7b-f**). Loss of neuronal PLA2G2F activity increased the average total abundance of ceramide (Cer), diacylglycerol (DG), lysophosphatidylethanolamine (LPE), phosphatidic acid (PA), and phosphatidylserine (PS) in the dentate gyrus (**Fig. 7d**). Almost all lipid species among those classes were upregulated in dentate gyrus of *Pla2g2f* cKO mice (**Extended Data Fig. 7g-k**). Differential expression analysis revealed significant changes in lipid species of triacylglycerol (TG), phosphatidylethanolamine (PE), cholesterol D7 (Ch-D7), monohexosylceramides (Hex1Cer), phosphatidylcholine (PC), and phosphatidylserine (PS) (**Fig. 4e,f**). Lipidome Identified Gene Enrichment Reactions (LIGER) were used to predict gene expression based on the presence and relative abundance of lipids in the samples. LIGER predicted multiple significant lipid conversion reactions that include PE and predicted associated genes orchestrating specific lipid reactions, including active conversion from O-PE to P-PE (PEDS1), suppressed conversion from O-DG to O-PE (PISD, PEMT) and from PS to PE and LPE (PISD, PLA2G4C). In addition, genes involved in the synthesis of phosphatidylserine (CDS1, PTDSS1/2) were predicted to be upregulated, and genes involved in the synthesis of PC (CHPT1, PISD) were predicted to be downregulated (**Fig. 7g**). LipidSig pathway analysis based on significantly different lipids (FDR<0.1) using Reactome (**Fig. 7h,** left) or KEGG (**Fig. 7h,** right) databases identified plasma lipoprotein assembly and clearance, membrane remodeling, glycerophospholipid biosynthesis, membrane-dependent receptor trafficking and lipid metabolism as potentially altered biological processes (**Supplementary Tables 4,5**). Corresponding genes included secreted phospholipase family members (*Pla2g2a/c/d/e/f*), regulators of lipid droplets (DGAT1/2), and lipoproteins (*ApoA/B/C/E*) whose human orthologs influence risk for cognitive decline and AD (eg: *ApoE2/3/4*)^75,76^.

Loss of *Pla2g2f* increased the relative abundance of neutral lipids like glycerolipids, including diacylglycerol and triacylglycerol, lipid species that are stored in organelles called lipid droplets ^89,90^ known to accumulate in microglia in dentate gyrus during aging and in AD^15,16,18,91,92^. Lipid droplet accumulation in microglia has been shown to impair phagocytosis and clearance of debris including myelin and amyloid plaques ^15,16,92^. To directly test if PLA2G2F mediates neuron-microglia signaling and regulation of lipid burden, we generated microglia-like cells (iMGLs) and excitatory neuron forebrain organoids from human induced pluripotent stem cells (**Fig. 5a**). To induce accumulation of lipid droplets in microglia, we treated iMGLs with the monounsaturated fatty acid, oleic acid (OA) ^18,93^. We then collected conditioned medium from excitatory neuron forebrain organoids expressing CamKIIα−*mCherry* or CamKIIα−*Pla2g2f*-T2A-*mCherry* and applied it to iMGLs overloaded with lipid droplets. Only the conditioned medium from forebrain organoids with *Pla2g2f* expression reduced lipid droplet accumulation in lipid overloaded iMGLs without changing iMGL area (**Fig. 5b**). Together, these data indicate that PLA2G2F is essential to maintain lipid homeostasis in the aged dentate gyrus and that PLA2G2F mediates neuronal-microglial signaling to alleviate lipid burden.

### Boosting *Pla2g2f* in an aging-sensitive AD model preserves memory and decreases amyloid burden

Aging is the biggest risk factor for AD and yet, mechanisms by which we can mitigate AD risk by directly modifying aging are poorly defined. Our data insofar suggests that PLA2G2F levels in the dentate gyrus dictate the extent of aging-associated pathophysiological changes and cognitive trajectory. Therefore, we tested whether boosting *Pla2g2f* expression in DGCs of an aging-sensitive mouse model of AD is beneficial. We selected the *App ^NL-F/NL-F^* knock-in mouse line in which the expression of mutant APP gene [Swedish (KM670/671NL) and Beyreuther/Iberian (I716F) mutations] is under control of its endogenous promoter, resulting in physiological (levels and spatial) expression of APP and overproduction of AbetaLJ42 and more faithful recapitulation of human AD pathological features ^94^. We injected lentiviruses expressing CamKIIα−*mCherry* or CamKIIα−*Pla2g2f*-T2A-*mCherry* into the dentate gyrus of 12 months old single amyloid precursor protein gene knock-in mice (*App ^NL-F/NL-F^*) and assessed cognitive performance in the pre-exposure-dependent contextual fear conditioning paradigm and quantified amyloid plaque number and size (**Fig. 6a**). Boosting *Pla2g2f* expression improved hippocampal dependent memory as evidenced by increased freezing in the fear conditioned context (**Fig. 6a,b**). Furthermore, we found a reduction in amyloid plaque size but not number in these mice (**Fig. 6c**).

## Discussion

Cognitive reserve theory posits the existence of compensatory resilience mechanisms that preserve brain health during aging ^41^, however, the identities of such mechanisms have remained elusive.

Our functional screen for synapse loss-induced factors, longitudinal analysis of *Pla2g2f* expression and mechanistic dissection of *Pla2g2f* functions during aging implicate this secreted phospholipase as a compensatory neuroprotective and cognitive resilience factor. Our data shows that PLA2G2F levels preserves lipid homeostasis to regulate the tempo of synapse loss, neuronal communication with microglia and astrocytes, neurogenesis, and memory impairment during aging. Loss of *Pla2g2f* in the dentate gyrus elevates lipid species associated with pathological aging, cognitive decline, and neurodegeneration as elaborated below. Boosting PLA2G2F levels improves cognition and reduces amyloid plaque burden in an aging-sensitive AD mouse model suggesting that efforts to decrease aging-associated pathological alterations such as inflammation, synapse loss and lipid dysregulation may delay onset of AD.

Studies in aged rodents, nonhuman primates and humans document perforant path degradation, loss of perforant path-DGC synapses and impaired dentate gyrus dependent cognitive operations in memory formation ^6–8,10,12^. We think that the elevation of *Pla2g2f* expression in sparse subsets of DGCs during aging is triggered by synapse loss and potentially by inflammation, DNA repair ^95^, and oxidative stress, known inducers of secreted phospholipase expression ^57,58^ and senescence ^96^. Preliminary analysis of *Pla2g2f* expression in human DGCs by single nucleus RNA sequencing revealed a similar highly sparse distribution within the dentate gyrus of cognitively intact aged individuals and to a lesser extent in aged individuals with Alzheimer’s Disease (Tosoni, Ayyildiz and Salta, personal communications).

The pattern of *Pla2g2f* expression observed in the dentate gyrus may reflect a heightened sensitivity of this subregion to aging-associated pathophysiological impairments ^6,9^. That said, we predict that *Pla2g2f* expression is also induced in other brain regions during aging to compensate for pathophysiological changes. Mapping the genetic and epigenetic landscape of *Pla2g2f* expressing cells will offer further insights into this thesis.

Our data supports a role for PLA2G2F functioning non-cell autonomously to preserve large diameter dendritic spines in DGCs during aging. This effect may be mediated through polyunsaturated fatty acids such as docosahexaenoic acid generated from hydrolysis of glycerophospholipids ^61,62^ or release of lysophospholipids to mediate intercellular communication in the neuronal milieu or blunting of levels of astrocyte-derived toxic long chain polyunsaturated fatty acid species ^26,97,98^. A non-cell autonomous role for PLA2G2F is consistent with how a sparse population of *Pla2g2f* expressing DGCs can influence dentate gyrus functions. Loss of *Pla2g2f* in the dentate gyrus resulted in elevation of phosphatidylserine (PS) and LIGER predicted multiple significant lipid conversion reactions leading to synthesis of PS (**Fig. 4f,g**). The elevation in PS may signify an increase in externalized PS that is an “eat me” signal for microglia to drive pathological synapse loss that we observe following *Pla2g2f* loss^99–101^. That PLA2G2F also contributes to large diameter dendritic spines by cell-autonomous mobilization of lipid species and continuous turnover of glycerophospholipids is also a possibility. Indeed, the elevation of ceramides seen following *Pla2g2f* loss is consistent with that seen in aging, MCI and AD and is associated with hippocampal atrophy ^102,103^.

Our lipidomics results showed significantly changed levels of many PE species and several potential conversion reactions involving PEs, suggesting a central role of PLA2G2F in PE metabolism, consistent with previous studies which showed a preference in the activity of PLA2G2F for the hydrolysis of PE or P-PE ^67^. Given that accumulation of PE and especially oxidized PE (O-PE) induced cellular ferroptosis^104^, a form of iron-dependent regulated cell death which is increased during aging and neurodegeneration^105^, our results suggests a potential role of PLA2G2F in protection against ferroptosis. Supporting this hypothesis, a previous study using genome-wide CRISPR activation screen identified PLA2G2F as a compensatory factor that inhibits ferroptosis^59^.

Lipid droplets are dynamic organelles increasingly implicated in the context of aging, APOE-associated risk for AD and neurodegeneration^15,16,91,17^. Typically consisting of neutral lipids at the core and polar lipids on the membrane, lipid droplets are essential to lipid storage and maintenance of balanced cellular lipid pool ^89^. However, accumulation of lipid droplets in cells such as microglia due to build up of neutral lipids like triacylglycerol and diacylglycerol, can compromise cellular processes such as phagocytosis and induce neuronal death^17^. PLA2G2F-dependent reduction of lipid droplets in iMGLs may restore functional competency of microglia in aging and AD and consequently, promote clearance of plaques and myelin debris. Given that cytoplasmic phospholipases regulate lipid droplet growth in a cell-autonomous manner^106^, it is likely that secreted phospholipases can regulate lipid droplets in other cells by hydrolyzing phospholipids in the extracellular space. Interestingly, many of the genes suggested by LIGER are regulators of lipid droplet growth. CDP-diacylglycerol synthase 1 (CDS1) has been shown to regulate the growth of lipid droplets in adipocytes; CDS1 knockdown led to the accumulation of phosphatidic acid (PA) and giant lipid droplets^107^. Choline/ethanolamine phosphotransferase 1 (CEPT1) and choline phosphotransferase 1 (CHPT1) are both enzymes which synthesize PCs, and CHPT1 knockout led to significantly increased cytosolic LD number and area, but decreased lipid droplet size^108^. Together, these data indicate that PLA2G2F is essential to maintain lipid homeostasis in the aged dentate gyrus and that PLA2G2F could mediate neuronal-microglial signaling to alleviate lipid burden.

Improved cognitive functions during aging may also restore peripheral physiology through hippocampal recruitment of subcortical circuits (lateral septum and hypothalamus) and induction of hypothalamic-derived hormones^109,110^. Consistent with this notion, boosting *Pla2g2f* expression significantly increased activity of aged mice during the night cycle, enhanced metabolic efficiency and prevented weight gain (Vicidomini, Yerevanian, Soukas and Sahay, unpublished observations).

Based on our findings, we propose that numerous cellular and synaptic changes are triggered to compensate for gradual loss of neuronal functions during aging and that potentiating some of these changes may be sufficient to confer cognitive resilience. Consistent with this framework, two other secreted phospholipase family members were recently shown to mediate compensatory roles in conferring resilience to high-fat diet induced inflammation-dependent weight gain^60^ and myelin damage from ischemia, respectively^61^. In conclusion, our studies suggest a critical role for dysregulation of lipid homeostasis during aging as driver of inflammation and synapse loss, two signatures of neurodegenerative disorders. This raises the possibility that restoration of lipid homeostasis by inducing or boosting expression of compensatory secreted phospholipases like PLA2G2F may harbor potential in modifying onset and trajectory of these disorders during aging.

## Methods

### Animals

All mice were group housed and experiments were conducted in accordance with procedures approved by the Institutional Animal Care and Use Committees at the Massachusetts General Hospital and NIH guidelines (IACUC 2011N000084). All mice were housed in a 12-h (7 a.m.–7 p.m.) light–dark colony room at 22°C–24°C with ad libitum access to food and water. Male C57Bl/6J, adult, middle-aged and aged mice (3,6,12,16,18 months) were ordered from National Institute of Aging (NIA). *Pla2g2f* ^f/f^ mice were generated previously ^67^ and obtained from Dr. Makoto Murakami. CamKIIα-Cre-ERT2 mice were previously generated ^111^and obtained from Dr. Richard L. Huganir. *App^NL-F/NL-F^* mice were generated as previously described ^94^and kindly donated to us by Dr. Takaomi C. Saido.

### Stereotactic Injection

Mice received carprofen (5mg/kg subcutaneously) before surgery and ketamine and xylazine(10mg/mL, 1.6mg/mL respectively) were used to anesthetize them.

Bilateral injections were performed using Hamilton microsyringes (Hamilton,NeurosSyringe7001) that were lowered to the target for 5 minutes prior injection, and injected at a rate of 0.1μL/min. The coordinates relative to bregma for the dentate gyrus subregion are: −2mm(AP), ±1.5mm (ML), −2.4(DV). The incision was sutured with coated vicryl sutures (EthiconUSLLC). Mice received daily injection of carprofen (5mg/kg, intraperitoneally) for three days after surgery.

### Tamoxifen Injection

Tamoxifen (10mg/mL, Sigma, T5648) was freshly prepared in 10% ethanol of corn oil (Sigma C8267). Mice were intraperitoneally injected for 5 consecutive days, once a day with a dose of 100mg/Kg.

### Dentate gyrus RNA sequencing and Bioinformatics

The dentate gyrus subregion was dissected from adult (3 months old) CaMKIIα rtTA; tetOKlf9/tetO-Klf9 mice in which *Klf9* expression was inducibly upregulated in the dentate gyrus following two weeks of 9-tert-butyl-doxycycline (9TBD) treatment as previously described ^31^. 9TBD treated tetOKlf9/tetO-Klf9 mice were used as controls. RNA extracted from dentate gyrus was subjected to polyA selection and library preps for next-generation Illumina Hi-seq (50bp, single-end reads, 25 million reads/sample). RNA sequencing read aligment was performed using bcbio-nextgen 1.0.2a0-a003647 (https://bcbionextgen.readthedocs.io/) against the GRCm38 / mm10 reference genome with transcript annotations corresponding to Ensembl release 84. Aligned counts generated by salmon 0.8.0 were used for downstream analysis. Differential expression analysis was performed in R 3.4.0 / Bioconductor 3.5 using bcbioRNASeq 0.0.18 and DESeq2 1.16.1. Analysis code is available upon request. Differentially expressed gene lists are provided in **Supplementary Tables 1,2.**

### Lipidomics

Mouse tissues were homogenized in phosphate-buffered saline using Bead Mill Homogenizer (VWR). Subsequently, lipids were extracted according to Folch’s Method. The organic phase of each sample, normalized by tissue weight, were then separated using ultra high-performance liquid chromatography coupled to tandem mass spectrometry (UHPLC-MSMS) method. UHPLC analysis was performed employing a C30 reverse-phase column (Thermo Acclaim C30, 2.1 x 250 mm, 3 μM, operated at 55° C; Thermo Fisher Scientific) connected to a Dionex UltiMate 3000 UHPLC system and a Q-Exactive Orbitrap high resolution mass spectrometer (Thermo Fisher Scientific) equipped with a heated electrospray ionization (HESI-II) probe. Extracted lipid samples were dissolved in 2:1 methanol:chloroform (v/v) and 5 μl of each sample was analyzed separately using positive and negative ionization modes, respectively. Mobile phase A consisted of 60:40 water/acetonitrile (v/v), 10 mM ammonium formate and 0.1% formic acid, and mobile phase B consisted of 90:10 isopropanol/acetonitrile (v/v), 10 mM ammonium formate and 0.1% formic acid. Lipids were separated over a 90 min gradient; during 0–7 minutes, elution starts with 40% B and increases to 55%; from 7 to 8 min, increases to 65% B; from 8 to 12 min, elution is maintained with 65% B; from 12 to 30 min, increase to 70% B; from 30 to 31 min, increase to 88% B; from 31 to 51 min, increase to 95% B; from 51 to 53 min, increase to 100% B; during 53 to 73 min, 100% B is maintained; from 73 to 73.1 min, solvent B was decreased to 40% and then maintained for another 16.9 min for column re-equilibration. The flow-rate for chromatographic separation was set to 0.2 mL/min. The column oven temperature was set at 55° C, and the temperature of the autosampler tray was set to 4° C. The spray voltage was set to 4.2 kV, and the heated capillary and the HESI were held at 320° C and 300° C, respectively. The S-lens RF level was set to 50, and the sheath and auxiliary gas were set to 35 and 3 units, respectively. These conditions were held constant for both positive and negative ionization mode acquisitions. External mass calibration was performed using the standard calibration mixture every 7 days. MS spectra of lipids were acquired in full-scan/data-dependent MS 2 mode. For the full-scan acquisition, the resolution was set to 70,000, the AGC target was 1e6, the maximum injection time was 50 msec, and the scan range was m/z = 133.4–2000. For data-dependent MS 2, the top 10 precursor selection in each full scan were isolated with a 1.0 Da window, fragmentation using stepped normalized collision energy of 15, 25, and 35 units, and analyzed at a resolution of 17,500 with an AGC target of 2e5 and a maximum injection time of 100 msec. The underfill ratio was set to 0. The selection of the top 10 precursors was subject to isotopic exclusion with a dynamic exclusion window of 5.0 sec. All data were processed using the LipidSearch version 5.0 SP (Thermo Fisher Scientific) and all lipid species with parent and at least one acyl chain detected (grade A, B) were reported. [grade A – all parent and acyl chains detected, grade B – parent and at least one acyl chain detected, grade C – no parent but both acyl chains detected, grade D – only single acyl chain detected]

#### Data Analysis

Statistical analysis was performed following a similar workflow described previously, the Comprehensive Lipidomic Automation Workflow (CLAW)^98,112–114^. Briefly, CLAW identifies differentially expressed lipids using a generalized linear model (GLM)^16^. The GLM uses a negative binomial distribution to model ion count data accounting for the technical and biological variability by incorporating a dispersion term using the common dispersion method ^115^. P-values were obtained through the likelihood ratio test and subsequently converted to the false discovery rate (FDR) adjusted for multiple testing using the Benjamini–Hochberg method. Lipids were considered significant based on a false discovery rate (FDR) value <0.1^116^. Lipid expression data was related to potential genes using LipidSig 2.0^117^ and the Chopra Lab in-house software, Lipidome Gene Enrichment Reactions (LIGER). LIGER predicts gene activity by matching lipids based on their subclass, chain length, and saturation using previously published statistical methods^118^. Building upon this initial work, we have expanded the database to include a more comprehensive range of lipid reactions to genes for pathway analysis. Additionally, methods were developed to address the high abundance of potential lipid matches, particularly between diacylglycerol (DG) and triacylglycerol (TG) lipid classes, thereby enhancing the robustness and accuracy of the lipid pathwayanalysis. By analyzing a variety of lipids with different chain lengths and saturations, LIGER accurately determines whether a gene reaction is generally upregulated or downregulated to assess how lipid molecules in sample relate to biological pathways. Code for all the data analysis is publicly available on: https://github.com/chopralab/PLA2G2F-Lipidomics. [Lipid name abbreviations: Cer – Ceramide, Ch – cholesterol, Ch-D7 – cholesterol D7, ChE – cholesteryl ester, DG – diacylglyceride, Hex1/2Cer – mono/dihexosylceramide, LPC – lysophosphatidylcholine, LPE – lysophosphatidylethanolamine, LPI – lysophosphatidylinositol, MG – monoacylglyceride, PA – phosphatidic acid, PC – phosphatidylcholine, PE – phosphatidylethanolamine, PI – phosphatidylinositol, PS – phosphatidylserine, SM – sphingomyelin, TG – triacylglyceride]

### iMGLs and human forebrain organoids generation

Human iPSCs were maintained at 37C and 5% CO2, in feeder-free conditions in mTeSR1 medium (Cat #85850; STEMCELL Technologies) on Matrigel-coated plates (Cat # 354277; Corning; hESC-Qualified Matrix). iPSCs were passaged at 60–80% confluence using ReLeSR (Cat# 05872; STEMCELL Technologies) and reseeded 1:6 onto Matrigel-coated plates. Dorsal forebrain spheroids were generated using a previously established protocol ^119^ with an adapted iPS seeding strategy as described previously in ^18^. Embryoid bodies (EBs) were generated using the same protocol as the spheroid induction protocol and seeded onto Matrigel-coated 6-well tissue culture plates at a density of 15-30 EBs per well. EBs were first differentiated into hematopoietic progenitor cells (HPCs) using the STEM diff Hematopoietic Kit (Cat#05310; STEMCELL Technologies). Following a previously established protocol, non-adherent HPCs were collected, centrifuged at 300 x g, and resuspended in 1 mL of microglia differentiation media (MDM) containing a mixed composition of half DMEM/F12 (Cat#11330-057; Thermo Fisher Scientific) and half Neurobasal media (Cat# 21103049; Gibco) supplemented with IL-34 (Cat#200-34; PeproTech) and m-CSF (Cat#300-25; PeproTech) ^120^. Cells were plated in 6-well tissue culture plates at 200,000 cells per well and maintained in MDM for at least two weeks before experiments. To induce lipid accumulation, 3 uM Oleic Acid (Cat# 03008; Sigma-Aldrich) was applied overnight to iMGLs. Live cells were imaged with an EVOS cell imaging system (Cat# AMF4300; Thermo Fisher Scientific). Images were processed in IMARIS (Oxford Instruments) to reconstruct cellular boundaries and quantify intracellular content as mean fluorescence intensity.

### Immunohistochemistry

Mice were anesthetized using intraperitoneal injections of ketamine and xylazine (10mg/mL, 1.6mg/mL respectively) and transcardially perfused with 4% paraformaldehyde in PBS. Brains were postfixed overnight in 4% paraformaldehyde at 4°C, then cryoprotected in 30% sucrose/PBS and stored at 4°C before freezing in OCT (OCT, FisherHealthCare) on dry ice. 35mm cryosections were obtained (Leica cryostat) and stored in PBS with 0.01% sodium azide at 4°C.

For immunostaining, floating sections were washed three times in PBS with 0.3% Triton X-100 (0.2% Triton X-100 for CD68-Iba1 staining) and then blocked in PBS with 0.3% Triton X-100 and 10% NDS (normal donkey serum) for 2 hours at room temperature. Incubation with primary antibodies was carried out O/N at 4°C. For 6E10 immunohistochemistry, sections were blocked in PBS with 0.3% Triton X-100, 5% NDS (normal donkey serum) and 5% BSA (Bovine Albumin Serum). Sections were washed three times with PBS for 15 minutes each time.

Fluorescent labeled coupled secondary antibodies (Jackson ImmunoResearch) were used at a final concentration of 1:500 in PBS/glycerol. Sections were mounted on glass slides and coverslipped with mounting medium containing DAPI (Fluoromount with DAPI, Southern Biotech).

List of antibodies used:

- Iba1:Wako Rb Chem 019-19741 1:500
- GFAP:DAKO Rb Z0334 1:500
- GFP: Aves Chicken GFP-1020 1:500
- CD68: Biorad Rat MCA1957 1:200
- RFP: Rockland Rb 600-401-379 1:200
- 6E10: Biolegend 80300 Mouse Anti-Ab 1-16 1:500
- C-FOS: SySy 226308 Guinea Pig 1:3000

### Viral constructs and virus preparation

Viral vectors were purchased from Vector Builder.

The following plasmids were used for viral deletion of *Pla2g2f*:

CamKIIα-*GFP* (Vector Name: pLV[Exp]-CaMKIIα_short>EGFP, vector ID: VB180531-1381ckb);

CamKIIα-Cre-T2A-*GFP* (Vector Name:pLV[Exp]-CaMKIIα_short>Cre(ns):T2A:EGFP, vector ID: VB180531-1380rte).

The following plasmids were used for virus mediated boosting of *Pla2g2f* expression:

CamKIIα-*mCherry* (Vector Name: pLV[Exp]-CamKIIα>mCherry, vector ID: VB170620-1070zsx);

CamKIIα-*Pla2g2f*-T2A-*mCherry* (Vector Name: pLV[Exp]-CamKIIα>>*mPla2g2f*[NM_012045.4](ns):T2A:mCherry, vector ID: VB170811-1047zxs).

Retrovirus preparation of RV-Tdtomato was performed as described previously ^31^. Lentiviruses were produced as described previously ^121^.

### Tissue collection and qRT-PCR analysis

cDNA preps were made from RNA isolated from dissected dentate gyrus. Total RNA was quantified using a NanoDrop spectrophotometer (Thermo Scientific) and then equal amounts of RNA were used for reverse transcription (SuperScript IV First-strand synthesis system, Invitrogen). qRT-PCR was carried out with SYBR green (BioRad) and primers with following sequences:

*Pla2g2f-*F 5’-AGCCTGGGTATGAAGAAATTC; *Pla2g2f-*R 5’-CAGTCTACCTCATCCATG;

*Gapdh*-F 5’-GCTTGTCATCAACGGGAAG; *Gapdh*-R 5’-TTGTCATATTTCTCGTGGTTCA.

### In Situ hybridization and RNA scope

ISH was performed using *Pla2g2f*-specific riboprobe generated from the coding sequence region of mouse *Pla2g2f* (NM 012045.4) corresponding to nucleotides 280-783 (exon 2-exon 5) as previously described ^31^. Color reaction was carried performed with nitro blue tetrazolium (NBT)/5-bromo-4-chloro-3-indolyl-phosphate (BCIP). Color reaction times were identical for every group. For quantification, three sections per mouse were analyzed using the mean intensity function in Fiji software. All images were captured using the same light intensity and exposure times. The mean intensity of the region of interest (minus the mean intensity of a selected background region) was averaged across images for each mouse and each group.

For RNA scope experiment fresh-frozen coronal serial sections (14um) were collected and RNA scope Multiplex Fluorescent Detection Kit was used to detect *Serpina3n* signal (Cat No. 430191 Probe Mm Serpina3n C1).

### Behavioral Procedures

Mice were handled for 3 days (~1 min per mouse/day) before behavioral experiments to habituate them to human handling and transportation from the vivarium to behavioral testing room. Before testing mice were transported in a holding room and allowed to habituate to the room for 30 min prior to behavioral testing. The videos for CFC were recorded and exported from Freezeframe (Actimetrics); videos for the other behavioral assays were analyzed with EthoVision XT 15 (Noldus).

### Open Field

The OF was a standard OF (41cmx41cm) as used in prior work ^121^.

Lux at very center of the OF was measured at 100. Each corner was measured to be 80 lux. Animals were tested for 10 or 15 minutes (as indicated in figure legends) before being returned to their temporary holding cages. The OF was cleaned in between each subject with 10% EtOH, and urine and feces was removed.

### Elevated Plus Maze

The EPM was a standard maze (two open arms (67 cm x 7 cm) perpendicular to two closed arms (67 × 7 × 17 cm) The four arms were separated by a neutral transition central square (5 × 5 cm). Lux levels: the center point of the EPM was 50 lux, the deepest points of the closed arms were 20 lux each, and both of the open arms were measured at 65 lux at their furthest ends. Animals were allowed to freely explore the EPM for 10 min before being gently removed and returned to their temporary holding cage. After testing, the EPM was cleaned with 10% EtOH, and any urine or feces were removed.

### Novel Object Location

An open field arena (41cm x 41cm) was modified as such: the outside of the clear plastic chamber was covered in white paper, up to the top of the chamber; the inside walls of the chamber was covered with white tape (up to ~15 cm from the bottom) to prevent the animals reflection in the plastic and improve contrast; on the inside of the chamber, a black vertical stripe (6 cm wide, rising to the top of the chamber; black duct tape) was placed in the middle of the northern wall; the plastic floor of the OF was covered in white tape, and then a thin even layer of bedding (~0.5 cm) was added throughout the entire chamber. Lux levels: 50 lux at the center, and 40 lux at each corner. This behavioral assay was adapted by a protocol previously described ^122^. Animals were first habituated to the chamber in the absence of any of the objects (3 days, 10 minutes session each day). Objects used were two identical clear Schifferdecker staining jars (5 cm x 7 cm x 9 cm); a pink piece of paper was inserted in each jar and the lid of the jars were secured with white puddy. The displaced object was counterbalanced such that the left or right object was equally displaced across groups. For training, the objects were placed in the chamber to the left and right of the black vertical stripe (3 cm from the edge of the object to the edge of the stripe), with the shorter end (7 cm) of the object being placed 0.5 cm away from the wall. For training, animals were allowed to freely explore each object in the arena for 15 minutes. For testing, either the left or right object was displaced to be directly opposite the black stripe, but now 3 cm from the southern wall (with the shorter end, 7 cm, of the object closest to the wall). For testing, a 10 min session was used. For scoring of objects exploration, EthoVision XT 15 was used to manually score the duration of sniffing behavior. Sniffing was considered as the nose-point of the animal facing the object (within 1 cm) with the body of the animal in the bedding. Standing or climbing on the object was not considered exploration. EthoVision XT 15 was also used for analyses of center-point movement in the arena.

### Novel Object Recognition

For Novel object recognition a different arena was used than that for NOL. Two new objects were also used. One set of objects (object A) were identical 150 mL glass bottles (body width: 5 cm circle; height: 12 cm; lid: 3 cm plastic circle), filled with tap water that was deeply darkened with blue food coloring. The other set of objects (object B) were two identical vertical coverglass staining jars (lid: 6 cm circle; base: 7 cm circle; body width: 4 cm x 4 cm square; height: 9 cm) filled with clear tap water. Animals were habituated to the empty arena for three days (10 minutes session each day). At training animals were allowed to explore either the A objects or the B objects (counterbalanced) for 15 min. At testing (24 hrs after training), either the left or right object from the day before (counterbalanced) was replaced with the other novel object. As for NOL, only sniffing was quantified as exploration of the object and was manually scored using EthoVision XT 15. Discrimination Ratio at training represents: (exploration time of left (L) object - exploration time of right (R) object) / (exploration time of left object + exploration time of right object). Discrimination Ratio at test represents : (exploration time of the novel (N) object - exploration time of familiar (F) object)/(exploration time of the novel object +exploration time of familiar object).

### Contextual Fear Conditioning

On day one mice were exposed to context A for 3 minutes in the absence of any aversive stimulus (foot shock). Features of context A were as follows: 10% acetic acid (no EtOH) was used to wipe down the chamber and for the chambers’ odor; a small houselight in the chamber was turned on (~50 lux; ~5 lux at grid floor); the room itself was dimly lit with small lamps; fans inside each cabinet were turned on to provide static background noise; the cabinet door was closed when the animals were inside; the test cage was equipped with gray metal walls and with a visible speaker; collection trays were placed underneath the grid; the shock sources were turned on. 24 hrs after context preexposure, mice underwent the same procedures as the day before; however, upon immediate entry into context A, mice received a single 2 sec, 0.75 mA footshock, and were immediately removed from the context. All animals were in the context no longer than 5 seconds. 24 hrs after the immediate shock mice were then tested for retrieval in context A (3 or 10 minutes as indicated in figure legends) in the absence of any shock in the same order as in the previous days. Freezing was automatically scored offline using FreezeFrame. Specifically, we utilized FreezeFrame’s trial viewer function to set the cutoff for freezing behavior. For each video, and while blind to group assignments, this threshold value was set at the bottom of the first trough of the motion index waveform generated by FreezeFrame (typically a value ranging from 5 to 50). Additionally, we set the minimum bout duration of freezing to be 1 sec or more.

### Image analysis of mossy fiber terminals, PV puncta, dendritic spines and microglia

Images were obtained from 3 sections per mouse hippocampus blind to treatment and genotype. A Leica SP8 confocal laser microscope and LAS software were used to capture images.

For MFTs, images were captured in the CA3ab subfield with a 0.3 μm step size of z-stacks using a 63x oil objective plus 5x digital zoom; for PV puncta, single confocal plane images were captured in the CA3ab subfields using a 63x oil objective plus 4x digital zoom. For dendritic spines, images were captured in the outer one-third of the molecular layer of the DG with a 0.3 μm step size of z-stacks using a 63x oil objective plus 4x digital zoom, then processed with the maximum-intensity projection using Fiji. For spine density, spines were counted manually for at least 20 μM of dendritic length per dendrite (20–30 dendritic segments were collected per mouse).

The Edge fitter plugin (www.ghoshlab.org) was used to measure head diameter (at the widest point of the spine head); 170 spines were analyzed per mouse to calculate spine size distribution.

For CD68 imaging, confocal z-stack images were captured in the granule cell layer of the dentate gyrus using a 40x water objective with a 1μm step size of z-stacks. Z stacks were flattened using the maximum intensity projection, and flattened images were quantified using Fiji. CD68 score was evaluated as previously described: 0 no/scarce expression; 1 punctate expression;

2 roughly one-third to two-thirds occupancy, and 3 greater than two-thirds occupancy for punctate expression or covering an entire cell or aggregated expression^123^. Microglial arbors were scored as follows: 0: thin long arbors with multiple branches, 1: thicker arbors with multiple branches, 2: thick retracted arbor with few branches, 3: no evident arborization, distinct amoeboid shape^123^.

### Quantification and statistical analysis

We adhered to accepted standards for rigorous study design and reporting to maximize the reproducibility and translational potential of our findings as described ^124^ and in Arrive Guidelines ^125^. All experimenters were blind to treatment conditions throughout data collection, scoring, and analyses. Statistical analyses were conducted using Prism v9 (GraphPad) and the minimum sample size was determined based on prior experience, existing literature, and a power analysis. Statistical significance was defined as p < 0.05. Analysis of variance (ANOVA) was used for multiple group comparisons. Repeated-measures ANOVA was used for comparison of groups across treatment, condition, or time. Detailed statistical analyses can be found in **Supplementary Table 6**.

## Author contributions

A.S conceived and administered the project, A.S and C.V co-developed the project, K.M.M performed synapse loss screen, C.V and T.D.G contributed to behavioral analysis and design, M.B.V and L.N performed the microglia-organoid co-culture experiments, C.V, L.E, M.L, M.R.F.S.S collected data, R.Y., C.B., and S.I. analyzed lipidomic data with G.C. provided supervision and funding, M.S, S.H.S, G.B and R.S performed bioinformatics, Z.W.L performed lipidomic profiling, T.C.W, L-H.T and A.S provided supervision, T.S, K.Y and M.M provided mouse lines, A.S provided funding, A.S and C.V wrote the manuscript with input from co-authors.

## Acknowledgements

We thank Sahay lab members, D.V.E, L.B.B.S and L.M.S.S for help with editing. A.S. acknowledges support from The Simons Collaboration on Plasticity and the Aging Brain, NIH R01MH111729, R01MH131652, R01MH111729-04S1, R01AG076612, R01AG076612-S1 diversity supplement, James and Audrey Foster MGH Research Scholar Award and MGH Department of Psychiatry. C.V was supported by The Simons Collaboration on Plasticity and the Aging Brain, MGH ECOR Fund for Medical Discovery (FMD) Fundamental Research Fellowship Award, a Harvard Brain Initiative Travel Grant, and a Brain & Behavior Research Foundation Young Investigator Award. T.D.G is a recipient of Brain & Behavior Research Foundation Young Investigator Award, a Harvard Brain Initiative Travel Grant, and a NIH K99/R00 Pathway to Independence Award (K99MH132768). M.R.F.S.S was supported by a HSCI summer research grant. L.-H.T acknowledges support from the Robert A. and Renee E. Belfer Family Foundation, the Cure Alzheimer’s Fund, The JPB Foundation, Joseph P. DiSabato and Nancy E. Sakamoto. M.M is supported by JSPS KAKENHI Grant Number JP20H05691 and AMED-CREST JP23gm1210013. K.Y is supported by JSPS KAKENHI Grant Number JP23K18289 and PRIME JP18gm5910012. Work by M.J.S and S.H.S was funded in part by the Harvard NeuroDiscovery Center and the Harvard Stem Cell Institute. Work by GC lab was supported, in part, by the United States Department of Defense USAMRAA award W81XWH2010665 through the Peer Reviewed Alzheimer’s Research Program, the National Institutes of Health (NIH) award, RF1MH128866 by National Institute of Mental Health, and NIH National Center for Advancing Translational Sciences U18TR004146 and ASPIRE Challenge and Reduction-to-Practice awards to G.C. Support, in part, from AnalytiXIN (Analytics Indiana) Fellowship in Life Sciences to G.C. and the Purdue Institute for Cancer Research funded by NIH grant P30 CA023168 is also acknowledged.

## Ethics Declarations

A.S and C.V are named co-inventors on U.S Patent PCT/US2023/01690, Vectors and methods for the treatment of neurodegeneration, delaying cognitive decline, and improving memory. G.C. is the Director of the Merck-Purdue Center funded by Merck Sharp & Dohme, a subsidiary of Merck and the co-founder of Meditati Inc. and BrainGnosis Inc. The remaining authors declare no competing interests.

## Data and Code availability

All data, code and materials used in this study are available in some form to any researcher for purposes of reproducing or extending the analysis. RNA sequencing data is accessible through GEO: GSE261906. Code for all our lipidomics data analysis is publicly available on: https://github.com/chopralab/PLA2G2F-Lipidomics. The lipidomics dataset is available upon request from the corresponding author.

## Supplementary information

**Extended Data Fig. 1.**
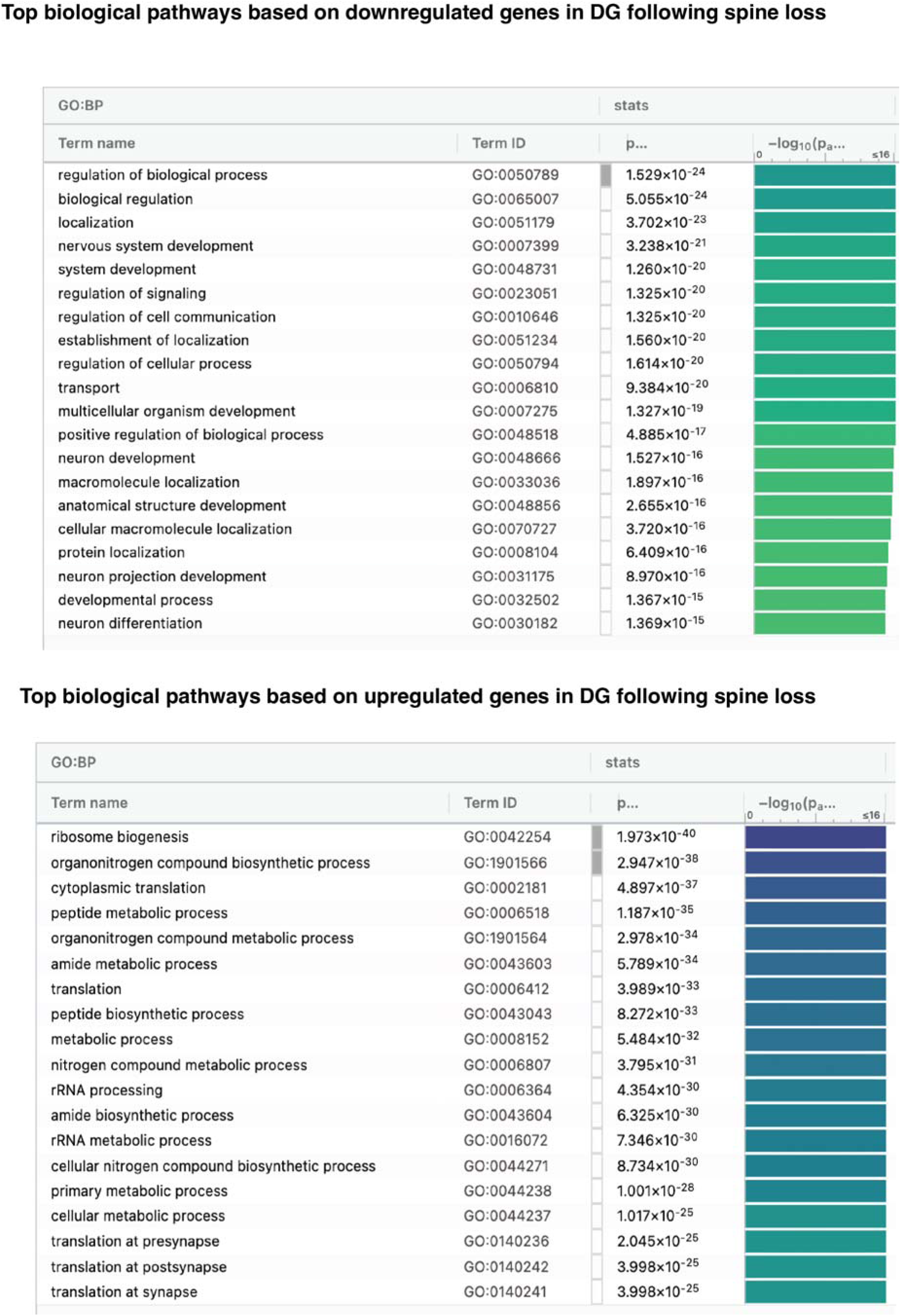
Down-and upregulated biological pathways in DGCs following inducible synapse loss. RNA was obtained from DGCs following genetic upregulation of Klf9 as described in Results and Methods. Functional enrichment analysis of RNA-seq dataset was performed using g:Profiler https://biit.cs.ut.ee/gprofiler/gost.

**Extended Data Fig. 2.**
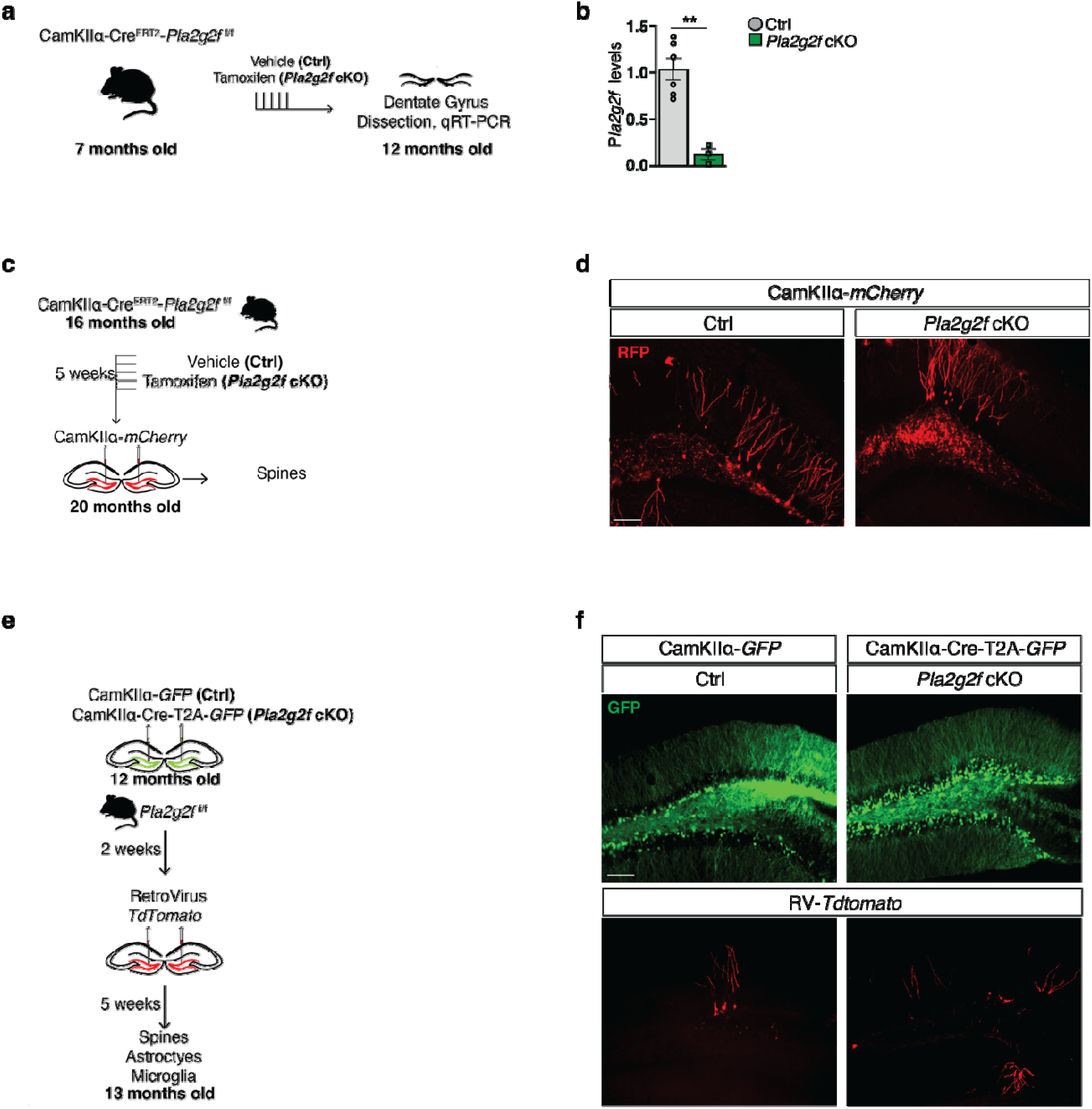
Supporting data for inducible deletion of *Pla2g2f* in DGCs. **a**, Experimental design to validate *Pla2g2f* deletion in CamKIIα-Cre^ERT2^-*Pla2g2f ^f/f^*bigenic mice injected with either vehicle (Ctrl) or tamoxifen (*Pla2g2f* cKO). Mice were injected at 7 months and 5 months later (12 months) dentate gyrus was dissected for RNA isolation and quantitative reverse transcription polymerase chain reaction (qRT-PCR). **b**, qRT-PCR of *Pla2g2f* expression in Ctrl and *Pla2g2f* cKO mice. Gene expression is measured relative to *Gapdh* (n=6 Ctrl, n=3 *Pla2g2f* cKO). Data are displayed as mean ± SEM; \*\**P*<0.005, [unpaired t-test] **c**, CamKIIα-Cre^ERT2^-*Pla2g2f ^f/f^* bigenic mice were injected with vehicle (Ctrl) or tamoxifen (*Pla2g2f* cKO), for *Pla2g2f* deletion, for 5 days; 5 weeks later, mice were injected with CamKIIα-*mCherry* lentivirus to label dendritic spines of DGCs; 4 months later spine analysis was performed. **d**, Representative images of DGCs of 20 months old mice expressing CamKIIα-*mCherry* lentivirus (scale bar 100μm). **e**, Experimental design to virally delete *Pla2g2f* in the dentate gyrus of 12 months old *Pla2g2f ^f/f^* mice and to label adult-born DGCs using retrovirus expressing *tdTomato*. Analysis of dendritic spine density of adult-born DGCs, astrocytes, and microglia was performed 5 weeks after retrovirus injection (13 months old mice). **f**, Representative images of dentate gyrus of 13 months old mice expressing CamKIIα-*GFP* (Ctrl) or CamKIIα-Cre^ERT2^-*GFP* and *tdTomato* expressing retrovirus to label adult-born DGCs (scale bar 100μm).

**Extended Data Fig. 3.**
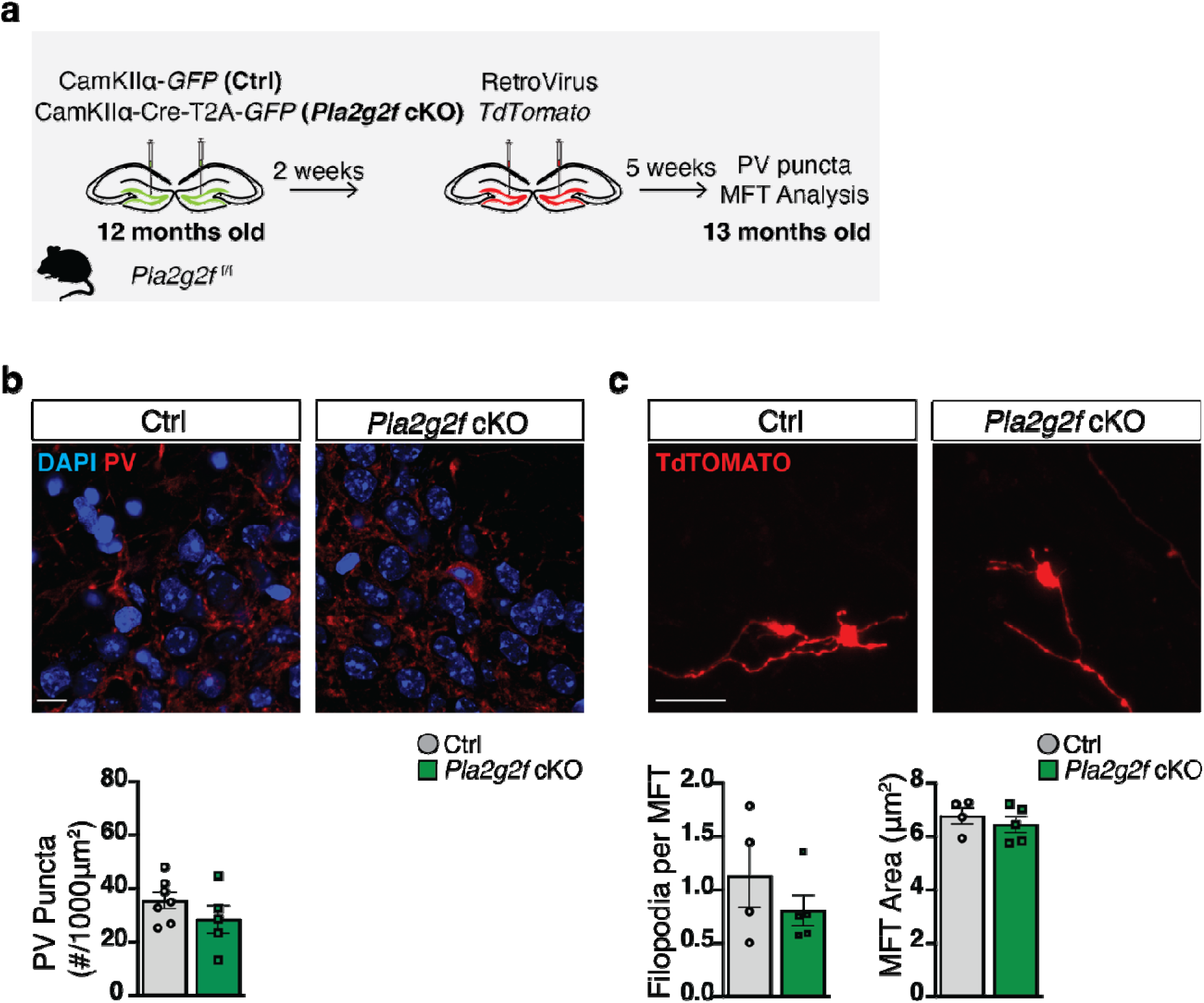
Deletion of *Pla2g2f* in DGCs of 12 months old mice does not change mossy fiber terminal size and mossy fiber terminal filopodial number of adult-born DGCs and number of parvalbumin inhibitory neuron synapses on CA3. **a**, Experimental design to virally delete *Pla2g2f* in the dentate gyrus of 12 months old *Pla2g2f* ^f/f^ mice and to label adult-born DGCs using retrovirus expressing *tdTomato*. Analysis of parvalbumin (PV) positive puncta density in subregion CA3ab and mossy fiber terminals of adult-born DGCs in CA3ab was performed 5 weeks after retrovirus injection (13 months old mice). **b**, Representative images and quantification of PV+ puncta density in CA3ab stratum pyramidale (n=7 Ctrl, n=5 *Pla2g2f* cKO; scale bar 10μm). Data are displayed as mean ± SEM **c**, Representative images and quantification of tdTomato positive filopodia per MFT (filopodial extensions) and mossy fiber terminal (MFT) area of 5 weeks old abDGC in CA3 (n=4 Ctrl, n=5 *Pla2g2f* cKO; scale bar 10μm). Data are displayed as mean ± SEM

**Extended Data Fig. 4.**
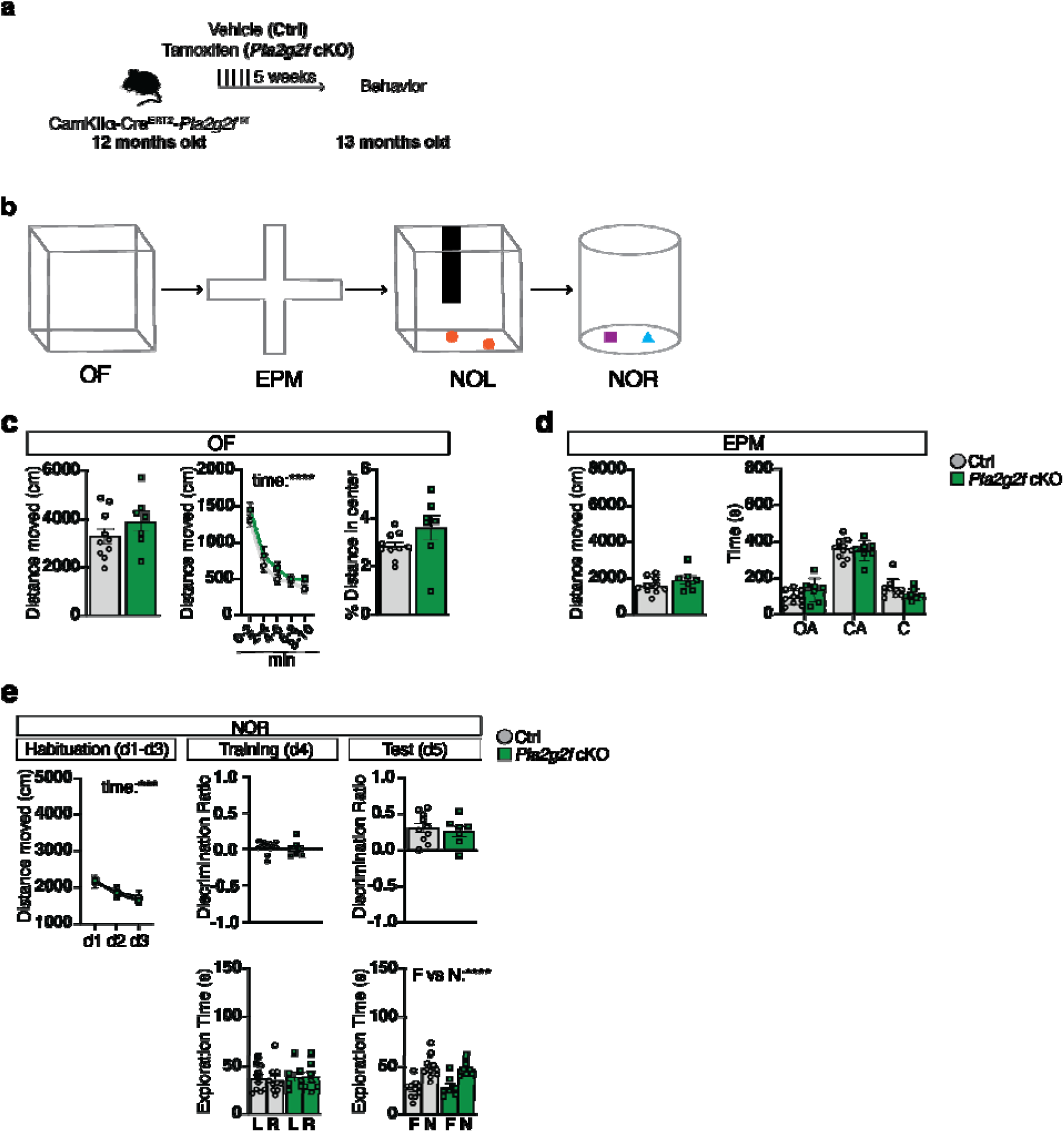
Deletion of *Pla2g2f* in DGCs of middle-aged mice does not affect anxiety-like behavior and novel object recognition. **a**, 12 months old CamKIIα-Cre^ERT2^-*Pla2g2f* ^f/f^ bigenic mice were injected with vehicle (Ctrl) or tamoxifen (*Pla2g2f* cKO), for *Pla2g2f* deletion for 5 days and behavioral testing was performed 5 weeks later in 13 months old mice. **b-e**, Ctrl and *Pla2g2f* cKO mice underwent behavioral testing for (**c**) locomotion (Open Field, OF), (**d**) anxiety-like behavior (Elevated Plus Maze, EPM) and (**e**) spatial learning and memory (Novel Object Location, NOL and Novel Object Recognition, NOR) (n=10 Ctrl, n=7 *Pla2g2f* cKO). Data are displayed as mean ± SEM;\*\*\*\**P*<0.0001, ***P=0.0010 [unpaired t-test in c (left and right panels), d, e (middle and right top panels); repeated measures two way ANOVA in c (middle panel) and e (top left panel, and middle and bottom right panel)].

**Extended Data Fig. 5.**
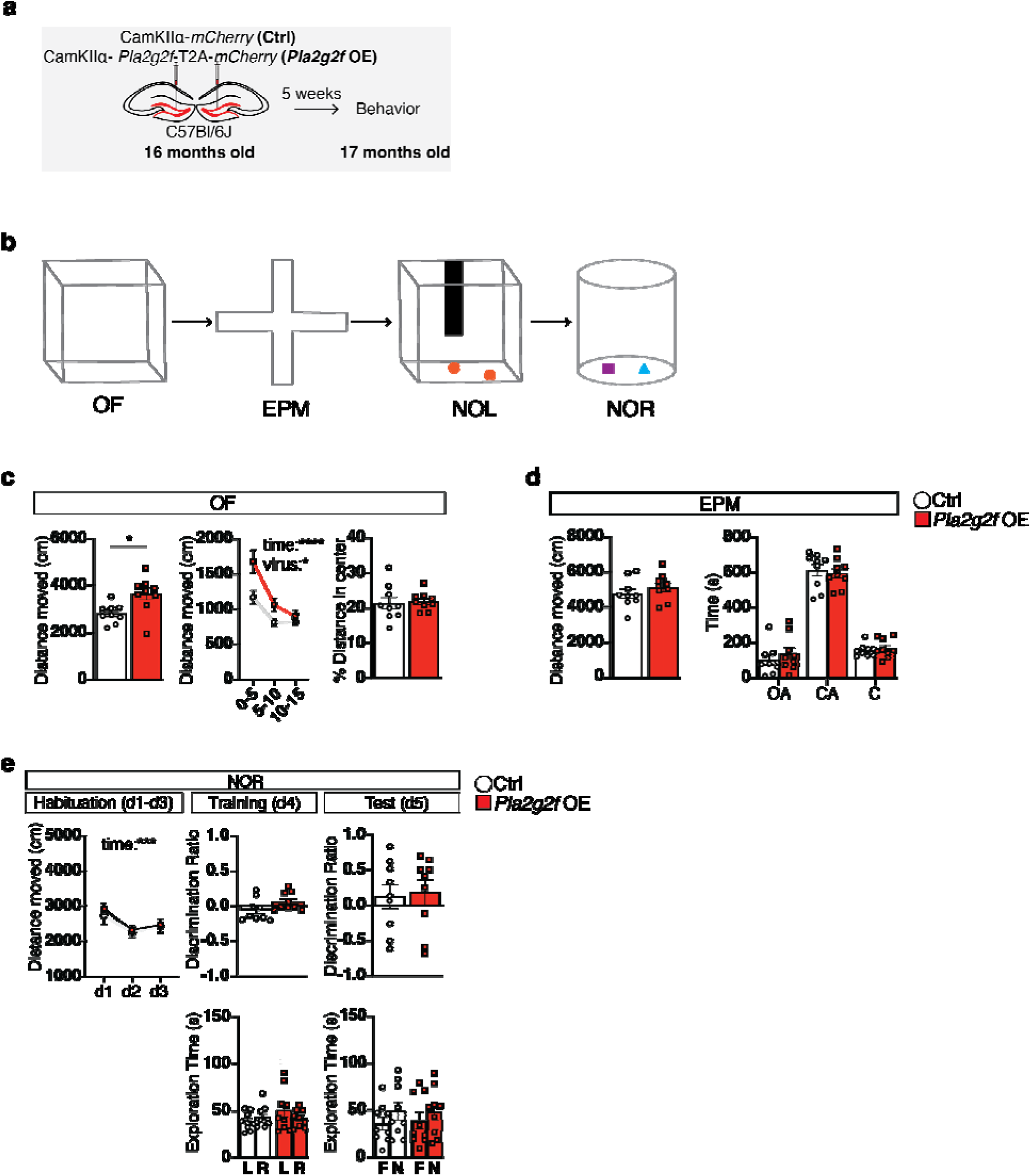
Boosting *Pla2g2f* expression in DGCs of aged mice does not affect anxiety-like behavior and novel object recognition. **a**, Experimental design to boost *Plag2f2* expression in DGCs of 16 months old of C57Bl/6J mice. Behavioral testing was performed 5 weeks following viral injections in 17 months old mice. **b-e**, Ctrl and *Pla2g2f* OE mice at 17 months underwent behavioral testing for (**c**) locomotion (Open Field, OF), (**d**) anxiety-like behavior (Elevated Plus Maze, EPM) and (**e**) spatial learning and memory (Novel Object Location, NOL and Novel Object Recognition, NOR). (n=9 Ctrl, n=9 *Pla2g2f* OE). For EPM, OA: Open arms, CA: closed arms. Data are displayed as mean ± SEM;\*\*\*\**P*<0.0001, \**P*<0.05 [unpaired t-test in c (left and right panels), d, e (middle and right top panels); repeated measures two way ANOVA in c (middle panel) and e (top left panel, and middle and bottom right panel)].

**Extended Data Fig. 6.**
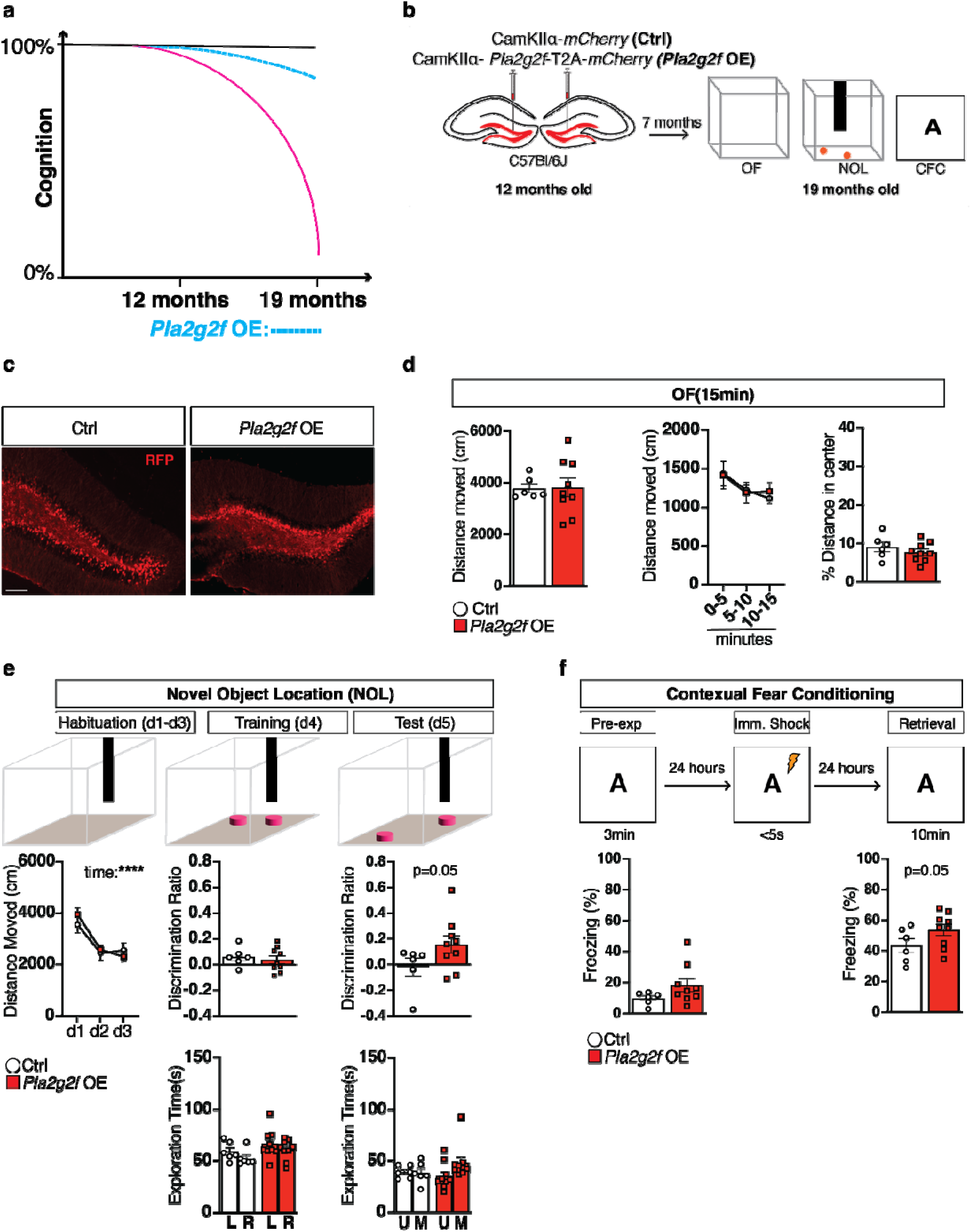
Boosting *Pla2g2f* expression in DGCs in middle-aged mice modifies trajectory of aging-associated cognitive decline. **a**, Schematic conveying the hypothesis tested in this experiment. Specifically, boosting *Pla2g2f* expression in middle-aged mice (12 months old) prevents age-associated cognitive decline in aged mice (19 months old). **b**, Experimental design to boost *Pla2g2f* expression in DGCs of 12 months old of C57Bl/6J mice. Behavioral testing (OF, NOL, CFC) was performed 7 months later (19 months old mice). **c**, Representative images of 19 months old mice dentate gyrus expressing either CamKIIα-*mCherry* (Ctrl) or CamKIIα-*Pla2g2f*-T2A-*mCherry* (*Pla2g2f* OE) lentiviruses (scale bar 100μm). **d**, Locomotion was assessed in Open Field (OF) behavioral test (n=6 Ctrl, n=9 *Pla2g2f* OE). Data are displayed as mean ± SEM. **e**, Hippocampal dependent learning and memory was assessed in Novel Object Location test. Discrimination Ratio at training represents: (exploration time of left (L) object - exploration time of right (R) object)/(exploration time of left object + exploration time of right object). Discrimination Ratio at test represents: (exploration time of the moved (M) object - exploration time of unmoved(U) object)/(exploration time of the moved object +exploration time of unmoved object) (n=6 Ctrl, n=9 *Pla2g2f* OE). Data are displayed as mean ± SEM. **f**, Contextual fear memory was assessed in pre-exposure-dependent contextual fear conditioning paradigm as the percentage of time spent freezing during retrieval. Baseline freezing was calculated during pre-exposure to context. (n=6 Ctrl, n=9 *Pla2g2f* OE). Data are displayed as mean ± SEM; [unpaired t-test in d (left and right panels), e (middle top panel) and f (left panel); unpaired one-tailed t-test in e (right top panel) and f (right panel); repeated measures two way ANOVA in d (middle panel) and e (top left and bottom middle and right panels)].

**Extended Data Fig. 7.**
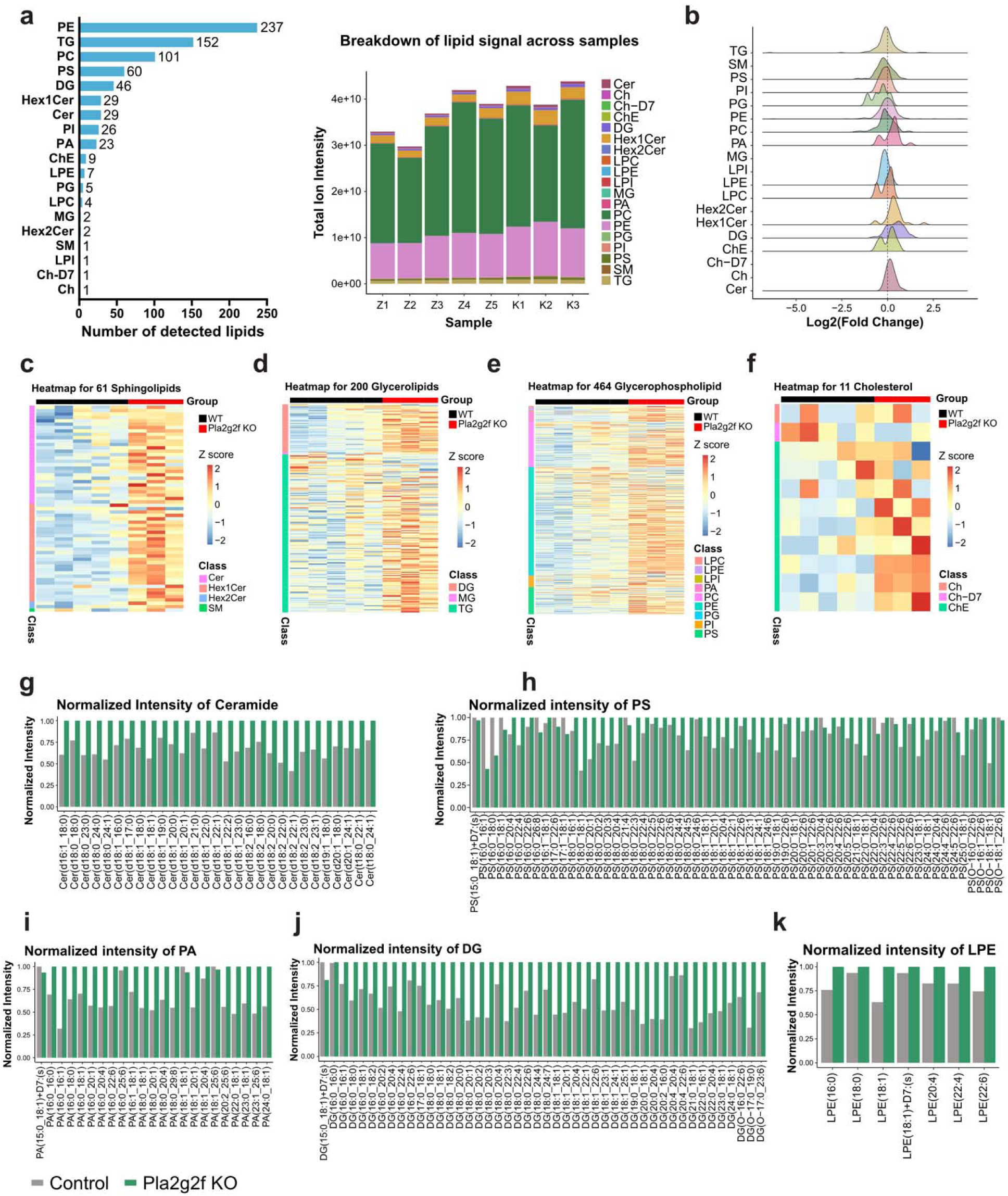
Supporting data documenting lipidomic changes in the dentate gyrus of *Pla2g2f* cKO mice. **a**, Breakdown of the number of lipid species detected in each class (left) and total lipid signal detected in each sample (right). **b**, Ridgeplot of average Log2(fold change) of lipids in Pla2g2f cKO mice vs. WT controls. Heatmap showing z score of lipid signals from c) sphingolipid, d) glycerolipid, e) glycerophospholipid, and f) cholesterol classes. Normalized ion intensities of lipids from g) ceramide (Cer), h) phosphatidylserine (PS), i) phosphatidic acid (PA), j) diacylglycerol (DAG), and k) Lysophosphatidylethanolamine (LPE) classes. Average lipid intensities were normalized by dividing over the maximal signal between the two groups.

**Supplementary Table 1.** List of downregulated genes in dentate gyrus following *Klf9* overexpression induced synapse loss. Analysis was done using a Benjamini Hochberg adjusted P value cutoff of 0.001 and a log fold-change (LFC) ratio cutoff of 0.25.

**Supplementary Table 2.** List of upregulated genes in dentate gyrus following *Klf9* overexpression induced synapse loss. Analysis was done using a Benjamini Hochberg adjusted P value cutoff of 0.001 and a log fold-change (LFC) ratio cutoff of 0.25.

**Supplementary Table 3. Lipidomics data from the dentate gyrus of Pla2g2f cKO and WT mice.** All data from UHPLC-MSMS were pre-processed using the LipidSearch version 5.0 SP (Thermo Fisher Scientific). Lipids with parent and at least one acyl chain detected (grade A and B) were analyzed using the EdgeR ^126^ package which calculates log2-transformed fold change (logFC), PValue, and false discovery rate (FDR) based on likelihood ratio test. Mean1 and Mean2 columns contain average ion intensity for Pla2g2f cKO group and WT group, respectively. The last 8 columns contain lipid ion intensity detected in each sample (Z: WT group, K: Pla2g2f cKO group).

**Supplementary Table 4. Reactome pathways enriched in Pla2g2f cKO vs WT group.** Lipid-related gene enrichment analysis was conducted using LipidSig 2.0^117^ and Reactome pathway database ^127^. Pathways were ranked by −Log10(p-value).

**Supplementary Table 5. KEGG pathways enriched in Pla2g2f cKO vs WT group.** Lipid-related gene enrichment analysis was conducted using LipidSig 2.0 ^117^ and KEGG database ^128^. Pathways were ranked by −Log10(p-value).

**Supplementary Table 6.** Complete statistics for all main and extended data figures.

